# Model-based whole-brain effective connectivity to study distributed cognition in health and disease

**DOI:** 10.1101/531830

**Authors:** Matthieu Gilson, Gorka Zamora-López, Vicente Pallarés, Mohit H Adhikari, Mario Senden, Adrià Tauste Campo, Dante Mantini, Maurizio Corbetta, Gustavo Deco, Andrea Insabato

## Abstract

Neuroimaging techniques are now widely used to study human cognition. The functional associations between brain areas have become a standard proxy to describe how cognitive processes are distributed across the brain network. Among the many analysis tools available, dynamic models of brain activity have been developed to overcome the limitations of measures like functional connectivity, via the estimation of directional interactions between brain areas. This opinion article provides an overview of our model-based whole-brain effective connectivity (MOU-EC) to analyze fMRI data, which is named so because our framework relies on the multivariate Ornstein-Uhlenbeck (MOU). We also discuss it with respect to other established methods. Once the model tuned, the directional MOU-EC estimate reflects the dynamical state of BOLD activity. For illustration purpose, we focus on two applications on task-evoked fMRI data. First, MOU-EC can be used to extract biomarkers for task-specific brain coordination. The multivariate nature of connectivity measures raises several challenges for whole-brain analysis, for which machine-learning tools present some advantages over statistical testing. Second, we show how to interpret changes in MOU-EC connections in a collective manner, bridging with network analysis. Our framework provides a comprehensive set of tools to study distributed cognition, as well as neuropathologies.

## 1 Introduction

The study of cognition has flourished in the recent decades due to the abundance of neuroimaging data that give access to brain activity in human subjects. Along the years, tools from various fields like machine learning and network theory have been brought to neuroimaging applications in order to analyze data. The corresponding tools have their own strengths, like predictability for machine learning. This article brings together recent studies based on the same whole-brain dynamic model in a unified pipeline, which is consistent from the model estimation to its analysis —in particular, the implications of the model assumptions can be evaluated at each step. This allows us to naturally combine concepts from several fields, in particular for predictability and interpretability of the data. We stress that our framework can be transposed to other dynamic models, while preserving the concepts underlying its design. In the following, we firstly review previous work on connectivity measures to set our formalism in context. After presenting the dynamic model (the multivariate Ornstein-Uhlenbeck process, or MOU), we discuss its optimization procedure to reproduce statistics of the fMRI/BOLD signals (spatio-temporal covariances), yielding a whole-brain effective connectivity estimate (MOU-EC). Then two MOU-EC based applications are examined: machine learning to extract biomarkers and network analysis to interpret the estimated connectivity weights in a collective manner. Meanwhile presenting details about our framework, we provide a critical comparison with previous studies to highlight similarities and differences. We illustrate MOU-EC capabilities in studying cognition in using a dataset where subjects were recorded in two conditions, watching a movie and a black screen (referred to as rest). We also note that the same tools can be used to examine cognitive alterations due to neuropathologies.

## 2 Connectivity measures for fMRI data

Among non-invasive techniques, functional magnetic resonance imaging (fMRI) has become a tool of choice to investigate how the brain activity is shaped when performing tasks [109, 98, 74, 31]. The blood-oxygen-level dependent (BOLD) signals recorded in fMRI measure the energy consumption of brain cells, reflecting modulations in neural activity [9, 48, 10, 106]. Since early fMRI analyses, a main focus has been on identifying with high spatial precision regions of interest (ROIs) in the brain that significantly activate or deactivate for specific tasks [32, 97, 152]. Because the measure of BOLD activity during task requires the quantification of a baseline, the brain activity for idle subjects became an object of study and developed as a proper line of research [17, 136]. This revealed stereotypical patterns of correlated activity between brain regions, leading to the definition of the functional connectivity or FC [22, 64]. Together with studies of the anatomical connectivity using structural MRI [139, 80], fMRI studies progressively shifted from focusing on specific ROIs to whole-brain analyses [39]. For example, high-order cognition involves changes in BOLD activity that are distributed across the whole brain [121, 29], which cannot be restrained to a small set of preselected ROIs. The typical description of the whole-brain activity in these methods is a ROI-ROI matrix, which we refer to as connectivity measure.

Recently, fMRI studies for both rest and tasks have also evolved to incorporate the temporal structure of BOLD signals in their analysis [83, 90, 30, 108, 69, 74, 25, 151], in addition to the spatial structure. Models have also been developed to formalize the link between the observed BOLD and the neuronal activity [61, 98, 41, 103, 69, 56, 151]. This has led to many definitions for connectivity measures and variations thereof. In this introductory section we present fundamental concepts about connectivity measures that set our model-based approach in context (see also the Glossary for definitions).

### 2.1 Model-based versus model-free approaches

A recurrent discussion opposes model-free and model-based approaches in neuroscience, which also applies for connectivity measures. For instance, FC calculated using Pearson correlation between time series (blue matrix in Fig. 1A) can be directly calculated from data —see Eq. (2) in Appendix D— and may thus seen as model-free. However, the interpretation of the FC values for the pairs of ROIs in the matrix is based on the theory of linear Gaussian models (LGMs), which assumes multivariate normal distributions for the data. The Pearson correlations is a connectivity measure that can be directly calculated on the generated activity. The underlying connectivity in the LGM, called the precision matrix, is closely related to the partial correlations (PC) represented by the green matrix in Fig. 1A. The PC symmetric matrix describes the undirected linear relationships between Gaussian variables that result in the observed correlations. Our viewpoint is thus that connectivity measures are always related to a model, so it is thus crucial to bear in mind their assumptions when interpreting their values. Note that other classifications of methods have been proposed and discussed in previous work, like directed functional connectivity measures versus effective connectivity measures [60], state-space models versus structural models for BOLD dynamics [144] and network-wise methods versus pairwise inference methods [16].

**Figure 1:**
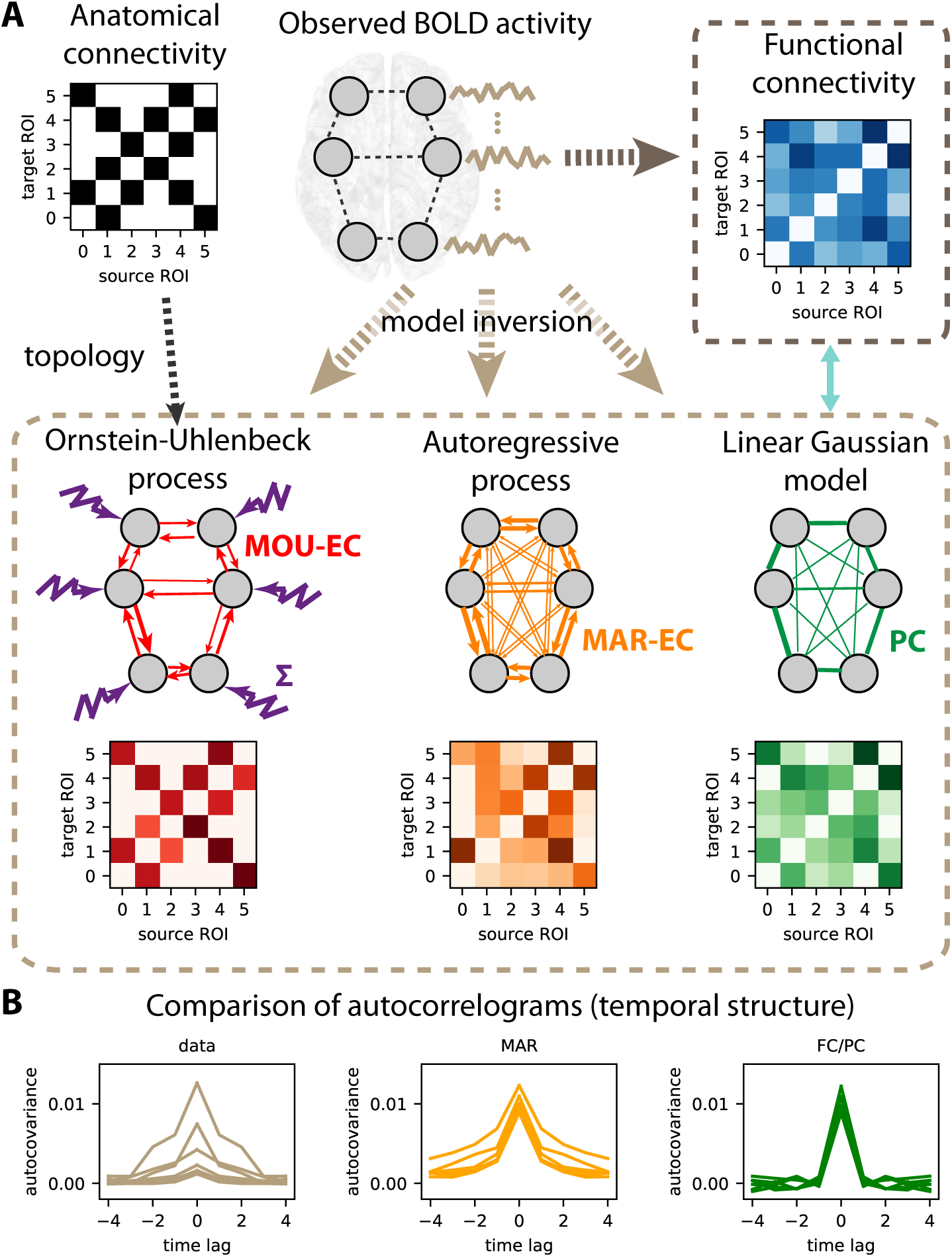
Connectivity measures. **A:** Schematic diagram of 6 brain regions of interest (ROIs), whose fMRI activity is captured by several types of connectivity measures. Here the activity is generated using a directed matrix. The corresponding structural connectivity (SC, black matrix) indicates the topology of anatomical connections between the 6 ROIs, which can be estimated using structural MRI. Functional connectivity (FC, blue matrix), here calculated using the Pearson correlation of the BOLD signals. The linear Gaussian model (LGM) assumes activity and corresponds to partial correlations (PC, green matrix). The multivariate autoregressive (MAR, orange matrix) assumes linear dynamics while taking the temporal structure of the data into account. Effective connectivity (EC, red matrix), which is the focus of the present article, depends on the choice for dynamic model, as well as input properties. Here the dark brown dashed box groups together the connectivity measures that involve a model inversion for their estimation, as compared to the light brown box that can be directly computed from the observed data. **B:** Autocorrelograms of the data (in gray, left plot) for 6 ROIs and two network models (middle and right plots). The linear Gaussian model related to the FC and PC (in green) has a flat autocorrelogram away from the central peak. In contrast, the profile of the MAR process (in orange) has a decaying exponential profile.

In the literature of whole-brain modeling, three families of connectivity measures have emerged:

- Structural connectivity (SC): It measures the density or probability of anatomical pathways that connect two ROIs, mostly via the white matter [129]. This led to the definition of the human connectome at the whole-brain level [139, 80].
- Functional connectivity (FC): Pairwise statistical dependencies between the observed activity of ROIs [22, 64]. Apart from the Pearson correlation of the BOLD signals [17, 136], other common FC measures include mutual information [86] and synchrony of instantaneous phases [24, 25]. Conceptually, FC corresponds a measure that can be applied to multiple time series, either the data or the model activity.
- Effective connectivity (EC): In this article we define EC as a measure of the directional relationships in a dynamic model. The original concept comes from electrophysiology [3], where EC determines how the stimulation of a neuron affects a target neuron (e.g. combining the source-to-target synaptic weight and the excitability of the target). It was then brought to neuroimaging in the 2000s when building models of the BOLD response and further developed in the context of the dynamic causal model (DCM), usually interpreted as neuronal coupling [61, 58, 144]. Note that it is also still used close to its original formulation when measuring stimulation-driven responses in neuroimaging [96].

To go beyond statistical relationships, the combination of several types of data requires a model, as with the SC and FC that are combined in the EC model (see Fig. 1A). Following, the choice of the model ingredients, especially their dynamics, has important implications that we detail in the following.

#### Glossary

- Generative model: Model of (dynamic) equations that generates a signal to be fit to empirical data. This definition is different from the definition in statistics where a generative model describes the joint probability distribution of observed and predicted variables (here the network activity), as opposed to a discriminative model that describes the conditional probability of the predicted variables with respect to the observed variables.
- Linear Gaussian model (LGM): Generative model of Gaussian variables with linear relationships. Its output variables have a flat autocovariance (apart from zero time lag) and is used to model noisy data without temporal structure.
- Multivariate autoregressive (MAR) process: Dynamic generative model in discrete time with linear relationships. Its output variables have both spatial and temporal correlations.
- Multivariate Ornstein-Uhlenbeck (MOU) process: Dynamic generative model in continuous time with linear relationships (referred to as connections). It is the equivalent of the MAR process in continuous time.
- Connections, interactions and links: In the main text connections refer to a direct and causal relationship between nodes (ROIs) in a dynamic model, whereas interactions are used to describe network effects that may be mediated by indirect pathways in the network (for example with FC or dynamic flow). Links are used in the context of machine learning as a common term for EC connections or FC interactions.
- Lyapunov optimization or natural gradient descent: Tuning procedure for the EC weights in the MOU network that fits the model FC covariance matrices to their empirical counterparts.
- Classification pipeline: Succession of machine-learning algorithms that aims to learn the mapping from input vectors of ‘features’ to output ‘labels’ (or categories). Here we use neuroimaging connectivity measures to predict cognitive conditions (like the task performed by a subject).
- Multinomial logistic regression (MLR): Linear multicategory classifier that assigns a coefficient to each input feature to predict the labels. It reduces to logistic regression for binary classification.
- k-nearest neighbor (kNN): Non-linear classifier that predicts the category of each new sample based on the most represented category over the k closest samples from the train set, given a distance metric or similarity measure.
- Biomarker: Subset of observed features (mainly EC/FC links) that enable a robust classification, often paired with weights as in the case of a linear classifier.
- Dynamic communicability: Measure of pairwise interactions between nodes (ROIs) in a network that takes indirect paths into account. In the present article, it corresponds to interactions over time for the MOU model.
- Dynamic flow: Extension of dynamic communicability that incorporates the effect of input properties in the MOU model.

### 2.2 Choice of model and interpretability

In our framework we borrow the EC terminology that can be understood in a broad sense as the directed connectivity in a generative model, here for BOLD activity. When using the LGM related to Pearson correlations taken as FC, one can take the partial correlation (PC, green matrix in Fig. 1A) as “LGM-EC”. As illustrated by the flat green autocorrelogram for non-zero time lags in Fig. 1B, the generated signals by the linear Gaussian model are independently and identically distributed (i.i.d.) variables and do not have temporal correlations. In contrast, the multivariate autoregressive process (MAR, in orange in Fig. 1A-B) has a directed connectivity (asymmetric matrix) and produces temporally correlated signals. When these models are fitted to data, they do not capture the same part of the data structure. When the MAR model is considered, the estimation results in directed connectivity that depends on the spatio-temporal correlation structure of the observed activity (or spatio-temporal FC). However, the linear Gaussian model does not “naturally” give directed connectivity when fitted to the spatial FC. Note that techniques based on optimization with regularization have been developed to enforce directionality in the estimated connectivity [128], though.

EC usually describes causal and directional relationships that interplay with other dynamic parameters in the model to determine the global network pattern of activity. When optimizing a model to reproduce BOLD signals, the parameter can be seen as representations or “projections” of the BOLD signals, in a top-down or data-driven approach [144]. Importantly, the estimated EC depends on the choice for the model dynamics. For example, the DCM was developed to formalize the link between neural and BOLD activities by explicitly modeling the hemodynamics [61, 140, 58]. The DCM-EC is thus the directed connectivity between the ROIs and determine their activities, from which the BOLD signals are generated via a hemodynamic response function.

We keep in mind that all interpretations of fMRI signals sit on the hypothesis that correlated BOLD activity between brain areas reflects the activity of their neuronal populations [73, 36, 100, 84], which in turn mediates the transmission of neuronal information [57]. However, many metabolic mechanisms like breathing (which are usually ignored in whole-brain models) alter the BOLD signals [116] and the adequate procedure to minimize the corruption of data and obtain a satisfactory signal-to-noise ratio is still under debate [115].

A distinct line of research [24, 41, 125, 118] has focuses on the development of whole-brain models in a more bottom-up fashion (model-driven), combining various datasets and biologically-inspired mechanisms, such as oscillators to produce rhythms observed in the brain. Such approaches allow for the study of the influence of specific parameters (as ingredients in the model) in collectively shaping the global network activity. Compared to connectivity estimation, a major distinction of many of those models is that SC is taken as the intracortical connectivity to focus on the influence of the local dynamics in shaping network activity patterns; see [103] for a review.

### 2.3 Consistency of pipeline from preprocessed data to modeling and analysis

The incentive for novelty has fostered the development of many methods in computational neuroscience. This makes results difficult to compare, especially for resting-state fMRI studies when no ground truth is available. Another caveat concerns network analysis when it is applied on connectivity estimates from time-varying signals (e.g. BOLD) with metrics (e.g. to detect communities) that do not genuinely relate to the physical values of the data. Our practical answer to this point is thus to develop for the same dynamic model a set of analysis tools, covering parameter inference [69] to machine learning and network analysis. Although the mathematical tractability of the model limits the richness of its dynamic repertoire compared to more elaborate bottom-up models [41, 125], it provides an intuitive understanding for the roles of parameters as well as analytical derivations for the parameter estimation or model interpretation. We believe that such an effort for formalization and consistency is beneficial to the field, both for didactic purpose and designing tailored methodology for complex models.

The present article follows the pipeline displayed in Fig. 2A. The second section presents the dynamic model and our recent optimization procedure [69, 66] to estimate MOU-EC for whole-brain fMRI data (Fig. 2B). It also discusses technical aspects in relation with other models used with neuroimaging data. The third section shows how connectivity measures can be used to predict cognitive states (Fig. 2C) and uncover their underlying structure. Importantly, the use of datasets with multiple tasks allows for an in-depth benchmarking of the connectivity measures. It also highlights some advantages of machine learning over statistical testing in making use of the multivariate nature of connectivity measures, transposing and abstracting concepts presented in a recent study for subject identification [112]. The fourth section bridges with network analysis [68, 67], interpreting changes in MOU-EC connections in a collective and model-based manner. In particular, it adapts community detection to our dynamic model (Fig. 2D).

**Figure 2:**
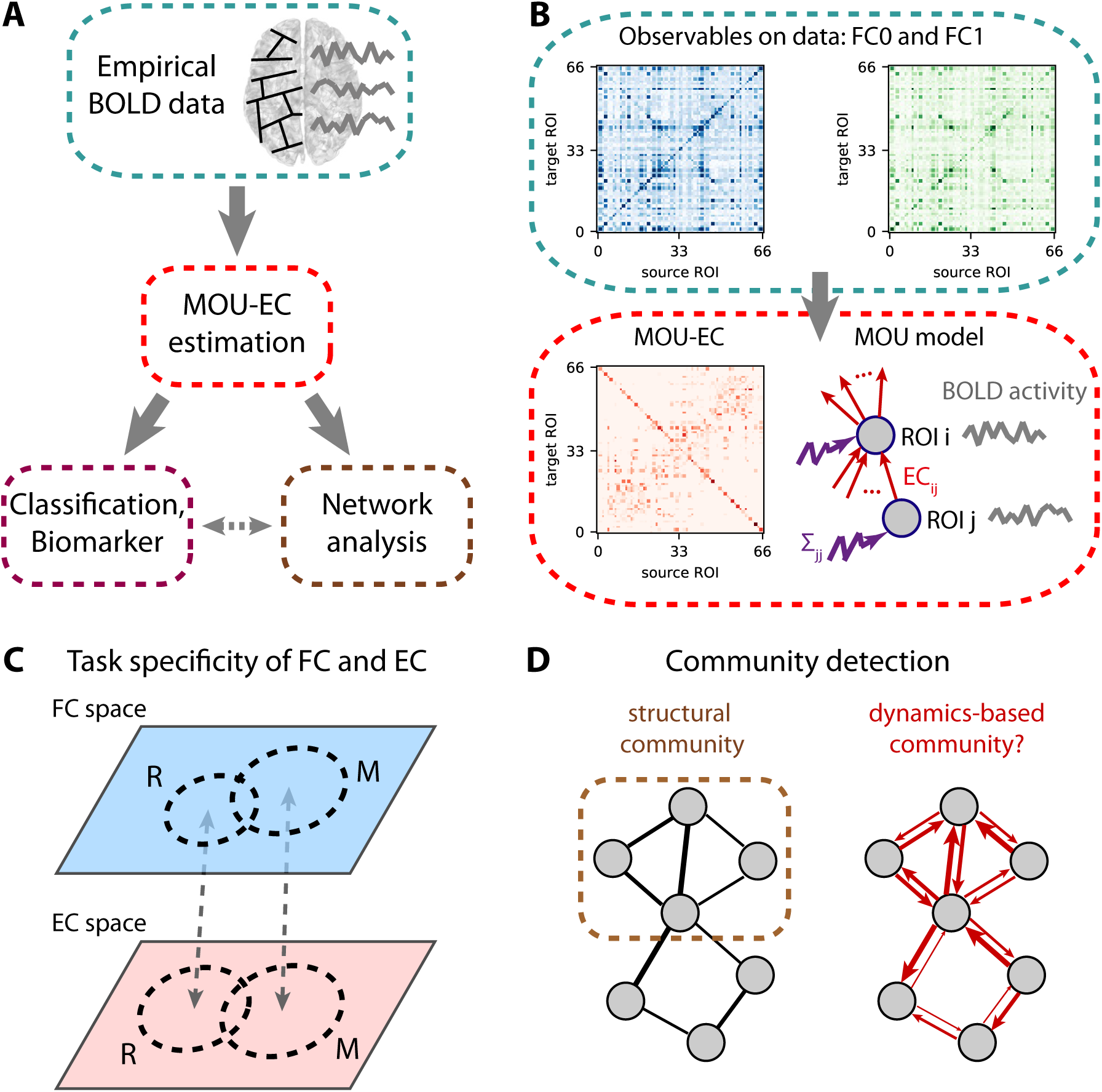
EC-based analysis of empirical BOLD signals. **A:** Pipeline for brain coordination analysis using MOU-EC. For each fMRI session, the model is optimized by tuning its parameters, especially its connectivity, to reproduce the statistics of the BOLD signals (depicted in the right side of the brain). A schematic parcellation is represented on the left hemisphere of brain. The estimated MOU-EC can then be used to predict cognitive states using classification algorithms. On the other hand, MOU-EC can be analyzed as a graph (or network) to uncover collective properties. A contribution of our approach is to link the network-oriented analysis with machine-learning techniques. **B:** From the BOLD signals, two FC matrices are calculated, FC0 without time lag in blue and FC1 with a time lag of 1 TR in green. The model parameters (MOU-EC and Σ) are optimized to reproduce these empirical FC matrices. Classification focuses on the model connectivity (e.g. MOU-EC), but the influence of Σ will be incorporated in the network analysis. See Fig. S1 for further details. **C:** Schematic representation of the FC and MOU-EC matrices (1 sample per session) in their own spaces for two cognitive states (R for rest and M for movie). For each connectivity measure, the prediction of R versus M is robust when the sample distributions represented by the dashed circles do not overlap. **D:** The left diagram represents community detection in a structural undirected network. The right diagram represents the MOU-EC networks, from which we want to define dynamics-based communities.

## 3 Capturing the whole-brain BOLD dynamics with MOU-EC

Using a generative model to reproduce the BOLD statistics, the optimized MOU-EC can be seen as an estimate that extracts spatio-temporal information about the BOLD *dynamics*. In fact, the main innovation of the modeling is the optimization procedure in Fig. 2B [69]. This section reviews important points about the underlying multivariate Ornstein-Uhlenbeck (MOU) dynamic model and its tuning. Further mathematical details can be found in Annex (see Fig. S1). Our approach aims to combine several key aspects:

- **Whole-brain** connectivity estimates are necessary to properly take into account the experimentally observed distributed information across distant regions [39, 29], without a-priori selection of ROIs.
- From the overall **spatio-temporal structure** of BOLD signals, the dynamic model only reproduces their covariances (without and with time lag, see Fig. 2B). This concerns the same fast timescale as recent studies [107, 108], which also demonstrated the influence of behavioral states such as sleep versus wake. The choice of BOLD covariances to be reproduced by the model is supported by previous results that showed that most of the information was in the second-order statistics of BOLD signals [86].
- **Causality** is inherent to the concept of EC [61, 58, 144] that are represented by the directed connections in the dynamic model (red arrows in Fig. 2B). From the estimated MOU-EC that best reproduces the data, we interpret strong MOU-EC weights as causal relationships and can examine their asymmetry to evaluate for a pair of ROIs which one drives the other. The optimization can deal with common inputs for ROIs, to explain the observed correlated activity by an interplay between the directed connectivity and correlated inputs [66].
- The MOU-EC **topology** is the inter-regional infrastructure, namely which connections exist and which do not in the brain network. When SC is available, its binarization (black matrix in Fig. 1A) can be used to constrain the MOU-EC topology in order to reduce the number of parameters to estimate. This enforces the model to “explain” changes in the FC by existing connections only, in contrast to PC and MAR (Fig. 1A). It is thus important to remember that MOU-EC estimates are not related to SC values.
- The optimization procedure tunes all MOU-EC weights while taking into account network effects. For each fMRI session, we obtain a **multivariate** estimate of more than 1000 parameters that represent the dynamical “state” of the brain activity. This contrasts with previous models that used the symmetric SC as connectivity matrix in dynamic models and focused on choosing or tuning the nodal dynamics with a few parameters only [41, 103, 125].
- A limitation of the model estimation procedure up to now is the assumption of stationarity for the BOLD signals over each fMRI session, which limits our approach to ongoing non-switching tasks.
- Another limitation of MOU-EC for the interpretation in terms of neuronal coupling is the absence of explicit hemodynamics in the model. This choice comes from the priority given so far to the estimation robustness (with simpler dynamics) over the biological interpretability, as will be discussed later.

### 3.1 Multivariate Ornstein-Uhlenbeck dynamics as generative model for BOLD signals

Formally, our model-based analysis is based on the multivariate Ornstein-Uhlenbeck (MOU) process that is described by Eq. 3 in Appendix C. It corresponds to a network with linear feedback that is the equivalent in continuous time of the discrete-time multivariate autoregressive (MAR) process. These dynamic systems with linear feedback have been widely used in neuroscience to model the propagation of fluctuating activity, mixing “noise” and structure, for example in modeling single neuronal dynamics [23], relating the connectivity and activity in a network [63] and defining neural complexity [143, 7]. It also corresponds to the linearization of nonlinear population models like Wilson-Cowan neurons [154].

The choice for the MOU dynamics is motivated by the balance between simplicity, which ensures tractable analytical calculation, and richness of the generated activity when modulating the parameters (especially the MOU-EC weights). Thus, it is well adapted to whole-brain data with parcellation involving many ROIs (≥ 100). In addition, the MOU dynamics implies exponential decaying autocovariances in the model, which have similar profiles to the empirical data (see the left plot in Fig. 1B corresponding to straight lines in log-plot in Fig. S1).

Intuitively, it can be understood as a network when fluctuating activity (akin to noise) is generated at each ROI and propagates via the recurrent EC. In other words, MOU-EC (red matrix in the bottom box) acts as a “transition matrix” and quantifies the propagation of fluctuating BOLD activity across ROIs. The MOU-EC matrix is usually sparse when its topology is constrained by SC (black matrix) to match anatomical white-matter connections between ROIs. The fluctuating activity for all ROIs is described by their (co)variance matrix Σ, which is diagonal in the present case (see the purple vector of variances). In a previous study, cross-correlations for Σ also involve common inputs to homotopic sensory ROIs [66].

### 3.2 Parameter estimation capturing network effects in BOLD propagation

To capture the BOLD dynamics (i.e. propagation of fluctuating activity), we use the two BOLD covariance matrices FC0 and FC1 in Fig. 2B, without and with time lag, respectively. This also ensures a one-to-one mapping between an FC configuration (a pair FC0 and FC1) and a MOU-EC configuration. This choice is in line with previous adjustments of DCM to model the resting-state that relied on the cross-spectrum, namely the Fourier transform of covariances with all possible time lags [58]. It is richer than fitting only an FC matrix without time lag [41, 103] and complies with a recent study of the task-dependent modification of the BOLD temporal structure at short time scales [108].

The MOU-EC estimate is obtained for the minimum model error in reproducing the empirical FC0 and FC1 for each session (see the fit plot between the boxes in Fig. S1). The optimization is akin to a ‘natural’ gradient descent [4] in that it takes into account the nonlinearity of the mapping from MOU-EC to the covariances FC0 and FC1. This arises because of the network feedback (even though linear) and may result in strong correlations (in FC) for disconnected ROIs (EC=0) provided strong indirect pathways connect them (via other ROIs). Appendix D provides the mathematical details of the optimization, which applies the gradient descent to both MOU-EC and Σ [66], extending the initial formulation with a heuristic optimization for Σ [69].

For illustration purpose, this article uses a whole-brain parcellation consisting with 66 ROIs. The SC density is 28%, giving 1180 MOU-EC weights to estimate (see Appendix A). Each fMRI session has 300 time points separated by TR = 2 seconds, so the number of data points is 66 × 300 ≃ 2.10^4^, about 16 times larger than the number of model parameters. EC should be like a summary of the BOLD signals: informative (not too short), but extracting and compressing relevant information (not too long). Our method was also successfully applied to the AAL parcellation with 116 ROIs and sessions of 5 minutes [112]. Typically, the MOU-EC estimation for a session and about 100 ROIs takes less than a minute of computation time on a desktop computer. For more refined parcellation or shorter sessions, the FC matrices may become quasi singular and the model estimates are expected to be noisier. The reduction of the number of parameters to estimate by using SC is crucial to work at the level of individual fMRI session and avoid overfitting. Here, overfitting would correspond to the situation where many distinct MOU-EC configurations in the model giving very similar pairs of FC0 and FC1 matrices, giving a larger overlap for MOU-EC than in Fig. 2C and the same small overlap for FC. The comparison between FC and MOU-EC as multivariate representation of cognitive states will be the focus of the next section. An important conceptual difference between MOU-EC and SC is that MOU-EC accounts for the dynamical properties of the brain activity, which are modulated when engaging a task. In other words, MOU-EC is hypothesized to account for the concentration of receptors or neurotransmitters, local excitability, etc., not only the density of the synaptic fibers.

### 3.3 Comparison with other approaches to extract information from BOLD signals

Other approaches have been proposed to characterize activity “states” based on the temporal structure of BOLD signals. For example, the ‘dynamic FC’ relies on sliding windows of several tens of TRs [90, 113, 117, 74], thereby focusing on changes in correlation patterns over minutes. Shorter timescales have been explored using instantaneous phases obtained using the Hilbert transform on the BOLD signals [25] or hidden Markov models (HMMs) [151, 18]. In contrast, the MOU-EC describes the BOLD propagation averaged over a session while assuming stationarity, as calculated in the corresponding statistics (covariances without lag and with a lag of 1 TR); Note that the “transition matrix” analogy for EC is at the level of the BOLD activity, not of hidden states as in the case of HMMs. Moreover, it does not involve a dynamic modulation of EC as used in the DCM [58, 98, 113].

A key innovation to tackle whole-brain fMRI data is the optimization constrained by SC that determines the network topology. This means the model has to explain the spatio-temporal FC structure using the existing connections only. The “prior” information related to SC avoids the preselection of ROIs and can be seen as an alternative to model comparison in choosing the best topology using structural equations [102, 92], Granger causality analysis [72] and early versions of the DCM [61, 144] —note that a recent DCM study incorporates SC for the network topology [137]. Likewise, the MOU-EC density usually depends on the threshold applied to SC and the best choice can be decided using model comparison, although the formulation may not be as natural as in the Bayesian framework.

Several fundamental properties were discussed a few years ago about the defining concepts of EC and DCM [144]. Beyond technical details, three main points are that DCM-EC corresponds to the connectivity weights in a dynamic model, that the model incorporates the hemodynamic response function (HRF) and that the estimation captures the BOLD dynamics, including the subsampling related to the low time resolution of BOLD signals. The EC terminology was borrowed from the DCM literature [58] because of the model-based aspect. The MOU-EC estimation was developed to solve the trade-off between robust estimation and application to large brain network (70+ ROIs) by using linear dynamics [69]. Since then, the DCM has been applied to whole-brain fMRI data [120, 56].

The FC0 and FC1 matrices in Fig. 2B embody the spatio-temporal BOLD structure in the range of “high” frequencies close to the Nyquist frequency equal to 0.5 Hz. The recent extension of the DCM to analyze resting-state fMRI data reproduces the BOLD statistics via the cross-spectrum [62, 120, 56], which is in line with our approach. Recall that this contrasts with earlier versions of the DCM that reproduced the BOLD time series themselves for stimulation protocols [61, 98]. Because it works in continuous time, the MOU model deals with the BOLD subsampling [69], unlike estimation methods relying on the discrete-time multivariate autoregressive process that may be sensitive to the subsampling of the BOLD signals [133]. Moreover, the observed lags between ROIs in FC —similar to cross-correlograms [107, 108]— are explained by the combination between the estimated inputs Σ and MOU-EC [65].

The time constant *τ*_*x*_ in the MOU model is identical for all ROIs in Eq. (3) in Appendix C, which corresponds to an abstraction of the HRF waveform response that was reported with a decay of 5 to 10 seconds [37]. An important difference of our model compared with the underlying dynamic model behind DCM [61, 144], as well as other bio-inspired network models [41], is the absence of an explicit HRF to link the neural activity to the measured BOLD signals [19, 59, 140]. Note that applications of the DCM to resting-state data involve a linearization of the HRF [120, 56]. Therefore, a foreseen extension of our approach is that of a state-space model with the MOU process generating the neuronal activity that is convolved with a linear HRF filtering [127]. The challenge is to keep the gradient-descent optimization tractable, as was previously done with Granger causality analysis [8, 52]. Incorporating the HRF may improve the reliability of the estimated EC [70, 111]. Note that an alternative consists in performing the deconvolution of the BOLD signals with respect to a HRF before analyzing the obtained neuronal signals, as was done with Granger causality analysis [36, 126, 153, 76] or other connectivity analysis [124].

## 4 Machine learning for extracting multivariate biomarker specific to cognitive states

Beyond their goodness of fit and how many “realistic” biological mechanism they might incorporate, models can be used to extract information from data in a top-down approach. Once tuned, the model parameters can indeed be used to discriminate cognitive conditions, as a representation of the observed data (BOLD signals here). This conceptually differs from finding the generative model of activity with the best fit. Nonetheless, it is expected that a model with poor fit hardly extracts any relevant information. Many recent studies have used connectivity measures to predict which tasks are performed by the subjects in the scanner [75], the pathological conditions of patients [95, 119] and individual identity [105, 54, 26, 5]. Machine learning is the standard for identifying biomarkers of neuropathologies [147] and is also widely used in voxel-based analysis for cognitive tasks [109, 152]. However, it is less frequently applied for cognition studies using connectivity measures [146]. For a given connectivity measure, a biomarker is a subset of connections (or links) that enables the robust identification of a category of fMRI sessions, e.g. a weighted sum of the matrix elements that exceeds a threshold for the MLR. In practice, changes in BOLD activity across cognitive conditions are mixed with individual traits and session-to-session variability. Disentangling these contributions is the key to obtain efficient biomarkers [112].

The present section addresses the following questions:

- How to cope with the multivariate nature of connectivity measures (e.g. > 1000 MOU-EC links)? We illustrate some advantages of machine learning compared to statistical testing, about multiple comparisons and beyond.
- What does classification tell about the underlying model? Linear and nonlinear classifiers apply different metrics on the estimated parameters, revealing the distribution of task-specific changes across brain regions.
- In a complex environment (e.g. many tasks), can we uncover a hierarchy in cognitive states (subgroups of tasks)?

Adopting the language from machine learning, ‘samples’ refer to fMRI sessions and the links of the connectivity measures are ‘features’ whose values are used to discriminate ‘labels’, which are the types of fMRI sessions here. To illustrate our framework, we use data that were previously analyzed [86, 99, 66, 44], whose details are summarized in Appendix A. Subjects in the scanner were recorded fixating a point in a black screen (two rest sessions R1 and R2) or a movie. The entire movie session was divided in three sessions (M1, M2 and M3), each corresponding to a different part of the movie. The vectorized MOU-EC estimates are displayed for all subjects and sessions in Fig. 3A. The question is then whether connectivity measures can capture information to identify the movie sessions individually. For example, the MOU-EC links indicated by the gray arrow exhibit strong changes between rest and movie (with small and large values, respectively).

**Figure 3:**
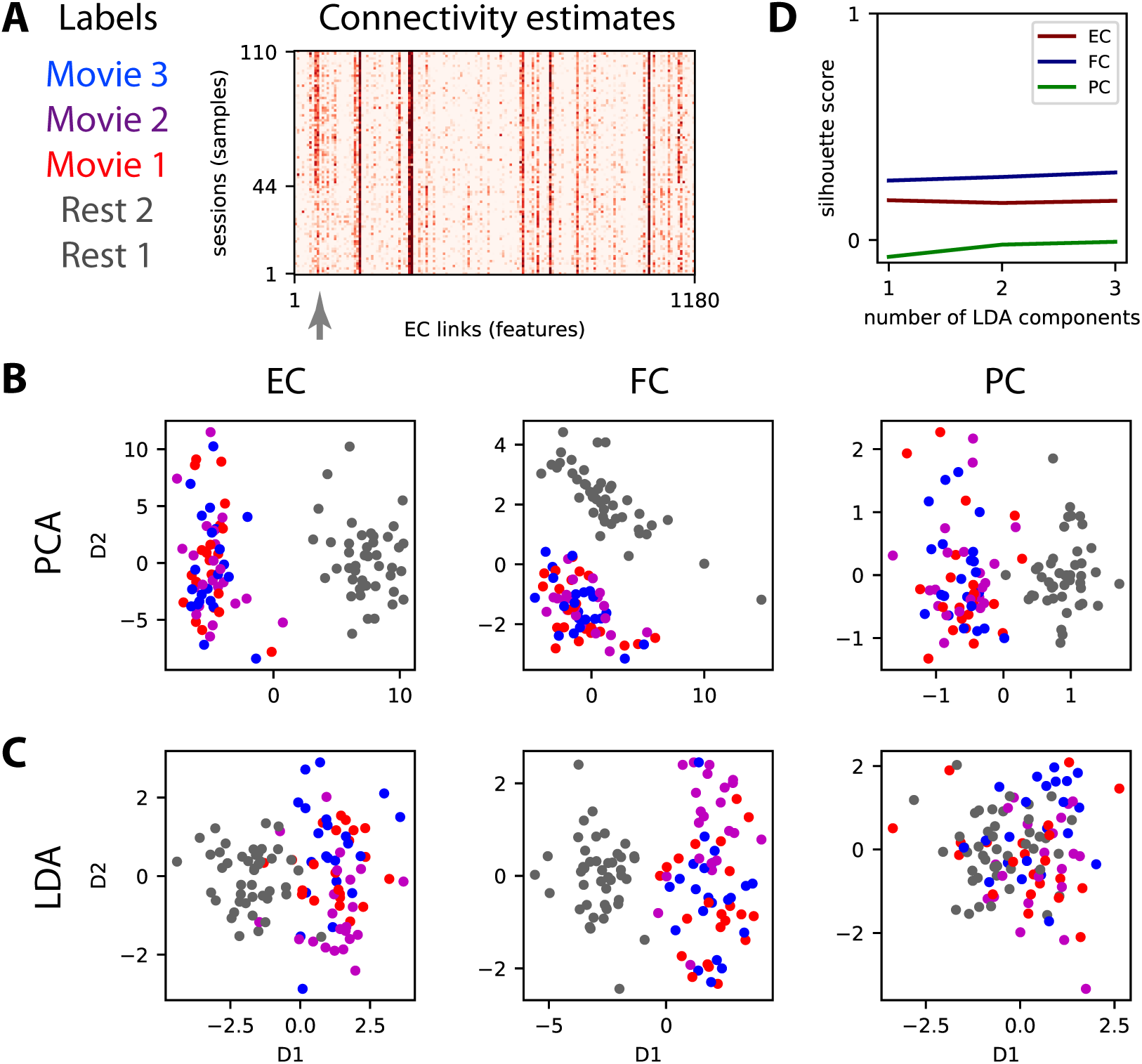
Dimensionality reduction of the connectivity measures. **A:** Connectivity estimates (here vectorized EC) for 5 sessions, 2 for rest and 3 for movie, for each of the 22 subject. The 3 movie sessions each correspond to a distinct part (10 minutes) of a movie watched and listened by the subjects. The gray arrow indicates connections (or links) that strongly differ between rest and movie. **B:** Principal component analysis (PCA) applied to the vectorized EC, FC and PC (one for each of the 110 sessions) where the same colors as in panel A correspond to the 4 tasks. For PCA the task labels are not used in the dimensionality reduction (unsupervised algorithm). **C:** Similar plot to panel B with linear discriminant analysis (LDA) for which the labels are used (supervised learning) to separate the 4 tasks for the 110 sessions. **D:** The silhouette coefficient measures the quality of the clustering for the 4 dot clouds when varying the number of LDA components (see x-axis).

### 4.1 Variability of connectivity measures with respect to cognitive labels

Before classification, the data structure can be explored to evaluate the similarity or distance across conditions between the connectivity measures, as representations of the BOLD signals. For this type of (passive) task, BOLD synchronization patterns during movie viewing between brain areas (e.g. visual and auditory) as reflected by inter-subject correlations [82] is expected to be also captured by the connectivity measures [66]. The gray arrow in Fig. 3A indicates some MOU-EC links with strong changes between rest and movie.

A usual technique for multivariate data is dimensionality reduction, for example using principal component analysis a.k.a. PCA [75] and independent component analysis a.k.a. ICA [27]. Here we compare two linear transformations, the unsupervised PCA in Fig. 3B with the supervised linear discriminant analysis (LDA) in Fig. 3C. The principal component is the direction in the high-dimension space with largest variability across the samples and successive components are ranked in decreasing order for their corresponding variability. It is called unsupervised because it does not use the task labels to compute the components, as can be seen for FC where the first component is not related to the task labels. In comparison, the components for LDA are ranked according to their variability *with respect to task labels*. This can be seen for FC where the first component for PCA does not reflect the separation between movie and rest (the second component does).

The separation of the tasks viewed using the connectivity measures can be evaluated using silhouette coefficients that measure the clustering quality [122], ranging from 1 for separated dot clouds to −1 for completely overlapping dot clouds. Applied to the LDA coordinates in Fig. 3D), we see a slight increase in silhouette coefficient when incorporating more components to separate the 4 clouds. This reflects the difficulty of the 4-task discrimination, as compared to that for the 2 tasks where a single component is sufficient [44]. The viewpoint here is that of clustering, assuming that the reduced space allows for isolating the clouds to characterize the cognitive states [75]. In the following, we rely on various classification techniques for the task discrimination.

### 4.2 Cross-validation for assessing the generalization capability of connectivity-based classification

Although machine-learning techniques are now increasingly used to analyze neuroimaging data, the generalization capabilities are not always properly verified [147]. For example, clustering algorithms applied on reduced-dimension components (see Fig. 3B-C) give a measure for the separation of the tasks viewed using the connectivity measures (Fig. 3D). Notice that all sessions are used for the dimensionality reduction in Fig. 3. This is a problem since the parameters tuning of the model can be influenced by the specific noise of the data samples and, in turn, the results will be not generalizable to new samples —a phenomenon called overfitting. To evaluate the robustness of a classifier, a cross-validation procedure is the standard for voxel-wise studies for activation maps [109, 152] and for clinical applications [119, 87]. This procedure consists in splitting the data samples in train and test sets that are respectively used to fit the classifier and to assess its performance, as described in Fig. 4A. Thus, one may have to include preprocessing steps in the cross-validation scheme to properly evaluate the generalization capability of the classification, for example with functional parcellations derived from the data that should be calculated on the train set only [21]. For LDA in Fig. 3C, this corresponds to setting a linear boundary using the dimensionality reduction on the train set and evaluating the performance on the test set. The data splitting can be repeated several times in order to assess the impact of different samples in the training and test set on the classification performance. Thus, the accuracy distribution provides an estimate of the generalization capability of the classification scheme to new samples. Different splitting strategies have been proposed and here we use the recommended one for neuroimaging applications is based on a ratio of 80%-20% for train and test sets, respectively [147]. We repeat the classification for 40 random splitting of the data. This is lower than the recommended 100 times and is justified by the rather low number of 22 subjects in our dataset, as explained in the box below together with further technical considerations about choosing a cross-validation scheme.

**Figure 4:**
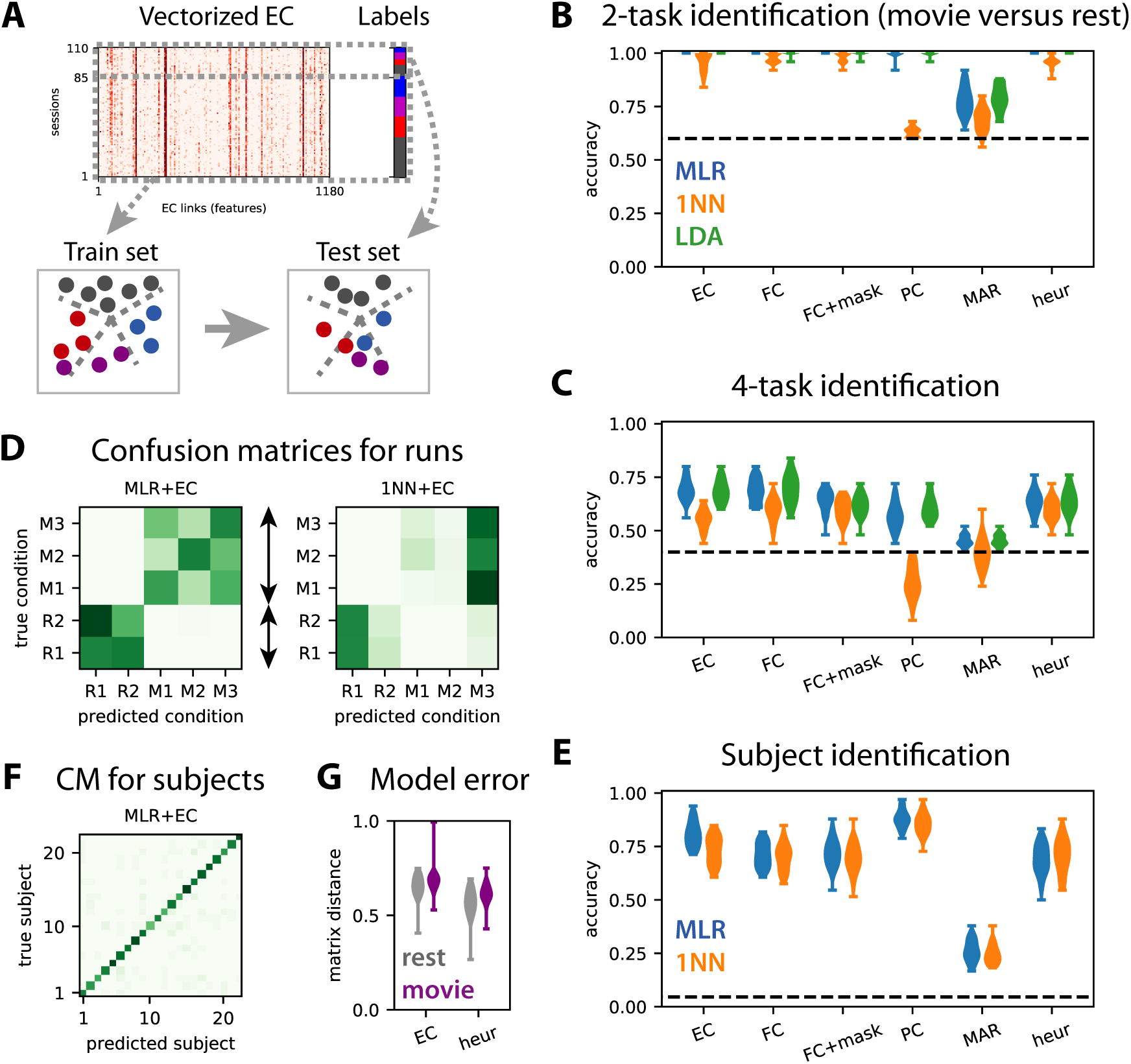
Train-test classification for task or subject identification. **A:** The train-test procedure consists in splitting the sessions in a train set and a test set. The train set is used to optimize the classifier (here find the best boundaries between labels, see dashed green lines), whose accuracy is then measured on the test set. **B:** Accuracy for the classification of test sessions to discriminate movie versus rest (2 labels) for the MLR, 1NN and LDA classifiers, as indicated by the colors. The connectivity measure are indicated on the x-axis: the masked FC is the subset of FC features corresponding to the SC mask used for the MOU-EC model; ‘heur’ corresponds to the symmetric connectivity matrix obtained using the heuristic optimization in Appendix D.2. The classifiers are trained with 80% of the subjects and tested on the remaining subjects, over 40 repetitions. The dashed line indicates the chance level, which corresponds to predicting always movie (3 sessions out of 5). The MLR gives perfect classification for all connectivity measures except for MAR. **C:** Same as panel B for the classification of session to identify the 4 tasks (3 movie sessions and rest). The chance level corresponds to predicting always rest (2 sessions out of 5). **D:** The left confusion matrix indicates the errors of the MLR classifier trained with 5 labels (2 labels for rest here), as indicated by the dark green off-diagonal pixels. As expected from panel B, rest and movie are well separated. The movie sessions (M1 to M3) are distinguishable to some extent, but not the rest sessions (R1 and R2). The right confusion matrix is the equivalent for the 1NN classifier. **E-F:** Same as panels C-D for the identification of the 22 subjects. Here we use 40 splits of the data are considered for 2 sessions in the train set and 3 sessions in the test set, irrespective of rest/movie and individually for each subject. **G:** Model error for the MOU-EC and heuristic optimizations in the two conditions.

#### Remark on cross-validation schemes

The recommended ratio for neuroimaging applications is 80%-20% for train and test sets, respectively, chosen by random draws and repeated about 100 times [147]. While we agree on this as a good general practice, we highlight that a slightly different scheme might be better suited when the number of available samples is rather low, as is often the case in cognitive studies like here with 22 subjects. Another issue concerns the independence or absence thereof of the measurements used as inputs for classification. For the example of Fig. 3, each subject is scanned in both resting-state and movie conditions, meaning that fMRI measurements are paired together. There, random resampling based on fMRI sessions without separating subjects in train and test sets would lead to more similar samples in the train and test sets than expected by chance. This would likely inflate the evaluated performance for generalization. A better solution is to apply the cross-validation to ‘groups’ (subjects in this case) instead of single samples (fMRI sessions). While it is possible to repeat random resampling of the groups several times, the chance of repeating the cross-validation for the exact same test set becomes non-negligible when the number of subjects is not sufficiently large. For our dataset, using 4 subjects for testing and the remaining 18 for training, there is almost 50% chance of getting the same split twice over 100 repetitions. The consequence would be an underestimation of performance variability. In such cases, an alternative valid option is the leave-one-group-out strategy, using a subject for testing and the remaining subjects for training, yielding a distribution of 22 accuracies. This procedure also has the advantage of highlighting individual differences in the dataset, showing for example if some subjects are easier or more difficult to predict. With our data, we find that repeating 80%-20% shuffle splits 40 times and performing leave-one-out for each of the 22 subjects leads to very similar accuracy distributions.

We consider task discrimination in two flavors: movie versus rest (2 tasks) in Fig. 4B and the movie sessions individually plus rest (4 tasks) in Fig. 4C. For each case, the performance evaluated using the test set should be considered relatively to perfect accuracy and chance level (dashed line). As expected from Fig. 3, the 4-task discrimination is more difficult that the 2-task one. The relative performance decreases by about a half of perfect accuracy for the 4 tasks compared to the 2 tasks. There are three interesting points coming from the comparison of the performances between the connectivity measures.

First, the MLR and LDA perform much better than the 1NN for the 4-task discrimination, both with MOU-EC and FC; this does not change when using kNN instead of 1NN. This agrees with previous results on subject identification [112] that are transposed to the present dataset in Fig. 4E. For task identification, the Pearson-correlation similarity for FC or MOU-EC does not seem the best choice, especially when the environment is complex. A technical point here is that LDA takes a long time to be trained for a large number of features or for many labels (subjects in Fig. 4E). Over all these results, the MLR seems the best practical choice for a classifier, as was found for clinical data [34].

Second, and more importantly, the masked FC performs worse than FC for the 4 tasks in Fig. 4C, demonstrating that the SC mask (with 25% density here) cannot be directly used to preselect the most informative FC links that contribute to the good performance. In contrast, MOU-EC has the same number of features as the masked FC and performs as well as FC, even though the silhouette coefficient in Fig. 3D is lower for MOU-EC than FC. Moreover, the MAR connectivity estimate, which extracts temporal information from BOLD signals as MOU-EC does, gives very poor accuracy, even for the 2-task classification. Partial correlations are somewhat in between. For subject identification, MOU-EC performs better than FC in Fig. 4E, in line with previous results [112]. Taken together, these results show that the dynamic model used for MOU-EC is a representation of the BOLD signals that extracts relevant information for both cognitive conditions and individual traits. In terms of dimensionality, MOU-EC can be seen as a more compressed version of the BOLD information than FC without loss of information.

Last, we consider the symmetric connectivity obtained by the heuristic optimization in Appendix D.2 that tunes the MOU model to reproduce the zero-lag correlation FC. In Fig. 4G the goodness of fit is slightly better for the heuristic method (smaller model error measure by the matrix distance) compared to EC. It is worth noting that the heuristic estimate leads to perfect accuracy for the “simple” classification rest versus movie (Fig. 4B). However, the accuracy decreases compared to MOU-EC for the 4-task and subject classifications by by 5% and 12%, respectively (Fig. 4C-E). This shows that the choice for the connectivity estimation method (with the corresponding measure on the BOLD activity) is crucial to efficiently extract information from the BOLD signals.

### 4.3 Capturing the hierarchy of cognitive states

The high dimensionality of connectivity measures allows for representing complex environments, such as the 4 tasks considered here. Beyond their ability to classify, the important question is whether MOU-EC or FC can capture the structure of the task categories —rest on one hand and the group of three movie sessions on the other. A similar goal was sought using clustering on FC to identify subtypes of depression [46], although concerns have been raised about the reproducibility of the results [45]. Clustering corresponds to unsupervised learning (without cognitive labels) and seeks a structure in the data for a given metric as with the first two components of PCA in Fig. 3B.

Instead, we rely on supervised learning and propose labels as an informed guess to a classifier, then assess whether they are meaningful using cross-validation. Beyond the performances of the 2-task and 4-task classifications in Fig. 4B-C, each classifier thus defines is a similarity metric for the connectivity measures. This is reflected in the confusion matrices in Fig. 4D, where the classification errors (dark off-diagonal green pixels) indicate which cognitive states are difficult to separate. By the naked eye, the structure of the left confusion matrix for the MLR determines a hierarchy that corresponds to the desired one, similar to the clustering in Fig. 3B and C. When asked to discriminate between the two rest sessions, the classifier gives a negative answer. The three movie sessions are identifiable even though the MLR classifier makes errors between them. In contrast, the right confusion matrix for the 1NN does not provide useful information. Similar matrices without a clear hierarchy are obtained for FC (not shown). This suggests that the Pearson correlation used as a similarity measure in the 1NN is not well suited to describe the cognitive states with sufficient flexibility. In other words, it is less the global EC/FC profile than specific links that significantly vary across tasks and should be the basis for building the task hierarchy. We do not enter into further details here, leaving the comparison between unsupervised and supervised learning in estimating label hierarchies in more depth for future work. As a last comparison, the confusion matrix for subjects in Fig. 4F shows that subjects are well identified and no particular structure in the errors is observed, indicating an absence of subject groups.

### 4.4 Biomarker formed of informative MOU-EC links: machine learning versus statistical testing

The general idea behind a biomarker is the indication of how and where to measure relevant information in the brain activity for the discrimination of several conditions, here related to cognition. The biomarkers for the 2-task and 4-task classifications in Fig. 4B-C are built by identifying the most informative links that contribute to correct classification, also referred to as support network. Practically, we employ recursive feature elimination (RFE) that efficiently works with the MLR classifier [78]. It provides a ranking of the features in order of importance and the support network consists of the first ranked links (with a cutoff when the classification performance based on those selected links reaches a maximum). A previous study compared these support networks of informative links for subjects and tasks to see whether the two classifications were biased one by the other [112]. Here we make another comparison with the two support networks in Fig. 5A, movie versus rest (9 black and 2 gray links) and for the 4 tasks (9 black and 35 red links). The 9 common links are also a hallmark of the hierarchy mentioned in the previous section, in the sense that the more 4-task biomarker extends the 2-task biomarker. Furthermore, the two RFE rankings are robustly aligned (Pearson correlation equal to 0.46 with p-value ≪ 10^−10^), as illustrated in Fig. 5C (left plot). If we compare with the support network for the FC links in Fig. 5B, we observe a similar overlap between the two support networks. Importantly, 5 FC links out of the 12 for the 2-task classification do not correspond to anatomical connections in SC, and 20 out of 30 for the 4-task classification. This suggests that FC changes reflect network-wise consequences rather than modulations of anatomical connections between ROIs (in dashed line). Nonetheless, we find that the RFE rankings for EC and FC are correlated, with a Pearson coefficient of 0.49 (and p-value ≪ 10^−10^), see the right plot in Fig. 5C.

**Figure 5:**
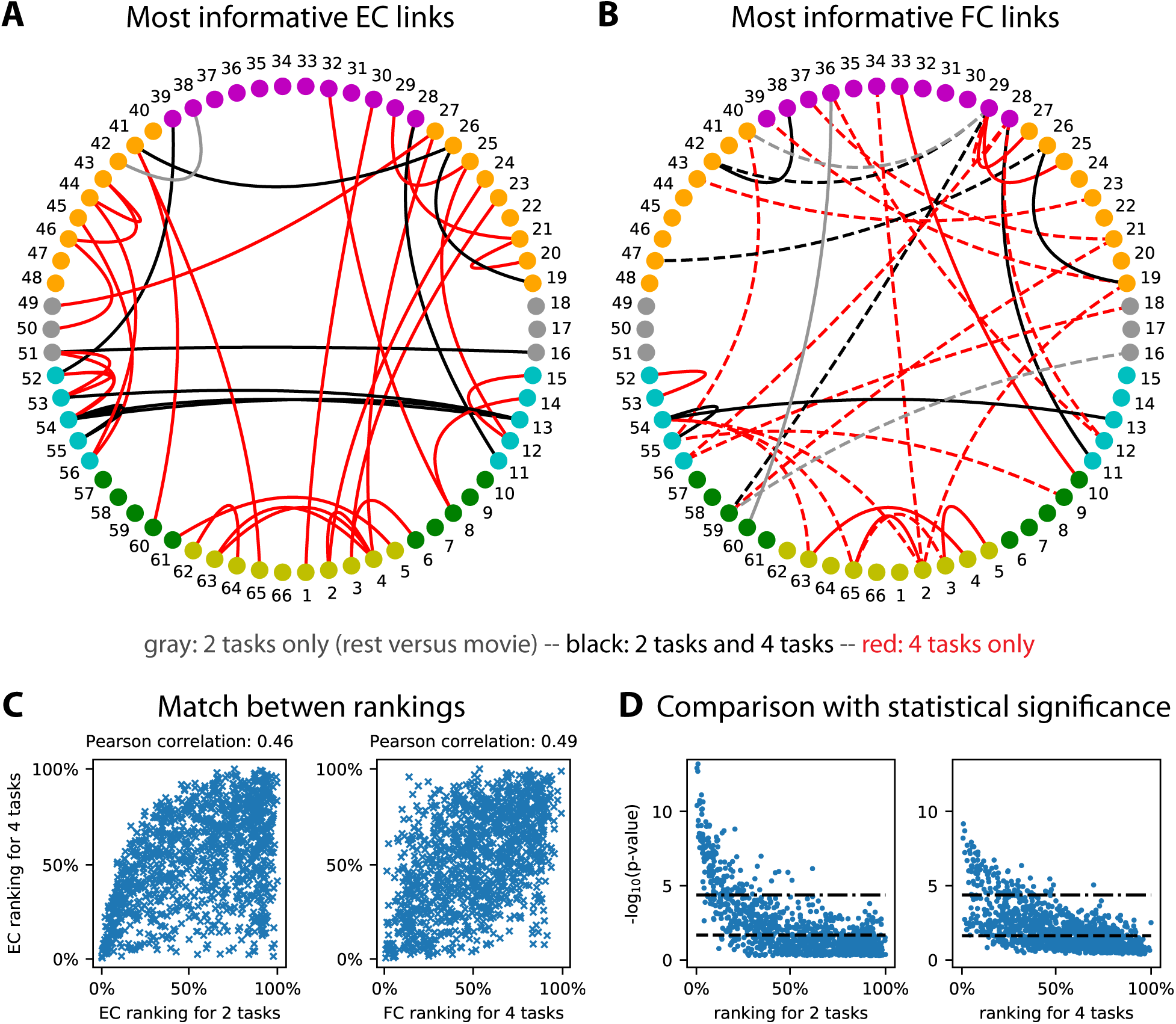
Biomarker for tasks and subjects. **A:** Support network of the 35 most informative MOU-EC links for the task identification, obtained using recursive feature elimination (RFE). The 10 black links are common to the easier discrimination between rest and movie, whereas the 29 red links are specific to the 4 tasks and the gray link is specific to the 2 tasks. The ROI colors indicate anatomical grouping: yellow = occipital, green = parietal, cyan = temporal, gray = central, orange = frontal and purple = cingulate. See also Table 1 for the ROI labels. **B:** Same as panel A for FC links. We obtain 8 common black links, 3 gray links for the 2 tasks only and 21 red links for the 4 tasks only. The links in dashed line correspond to ROI that are not connected in SC. **C:** Match between EC rankings for the 2 and 4 tasks (left plot), as well as between EC and FC rankings for existing EC links. The Pearson correlation between the matched rankings is indicated above. **D:** Correspondence between p-values for Mann-Whitney tests and RFE rankings for the 1180 MOU-EC links. For the 2 tasks, we show two significance thresholds for multiple comparisons: the Benjamini-Hochberg correction (dashed line) and Bonferroni correction equal to - log_10_(0.05*/*1180) (dashed-dotted line). For the 4 tasks, the maximum of the p-values for the 6 pairwise comparisons is plotted, with the same Bonferroni threshold.

Now we discuss the pros and cons of machine learning compared to statistical testing in finding correlates (i.e. link changes) of cognitive tasks. Formally, these two methods correspond to distinct questions that relate to opposite mappings from measured variables to conditions: changes in the variables across conditions for statistical testing and the prediction of conditions based on the variables for machine learning. However, they can be used to address the same practical question, especially when interpreting data in two conditions (as here with rest versus movie). In neuroimaging data a crucial point relates to the high dimensionality of connectivity measures, which leads to repeated hypothesis testing —are MOU-EC values different across two conditions?— over the 1180 connections. This calls for correction for multiple comparisons, which is problematic as the number of connections scales with the square of the number of ROIs in general. We illustrate this point by calculating the p-values for the non-parametric Mann-Whitney test between rest and movie MOU-EC weights for each connection (left plot in Fig. 5D). To identify links that strongly differ across conditions, we use the significance threshold with Bonferroni correction (dashed-dotted line), here − log_10_(0.05*/*1180) for 1180 MOU-EC links, or better with Benjamini-Hochberg correction [11] (dashed line). The same can be done for the 4 tasks (right plot), which involves 6 pairwise tests between the 4 cognitive states. For each connection we retain the smallest p-value among the 6 (as it is informative for at least a pairwise discrimination). For the Bonferroni correction, this gives 98 connections passing the significance threshold for the 4-task, to be compared with 153 for 2 tasks. Using Benjamini-Hochberg correction, we find for the two classifications 554 and 498 links, respectively. This does not satisfactorily reflect the increase of complexity of the cognitive labels —more labels require more connections in the biomarker— while the support network obtained using RFE in Fig. 5A provides a satisfactory description. More importantly, the comparison of the mapping between p-values and the RFE ranking in Fig. 5D also illustrates that efficient classification may rely on link with poor p-values, and that this phenomenon may be stronger when the number of categories increases (like labels here). This conclusion also holds when using the parametric Welch t-test instead of the non-parametric Mann-Whitney test. Beyond the details of the present example, machine learning seems better suited than statistical testing to deal with connectivity measures in a context of multiple labels.

**Table 1:**
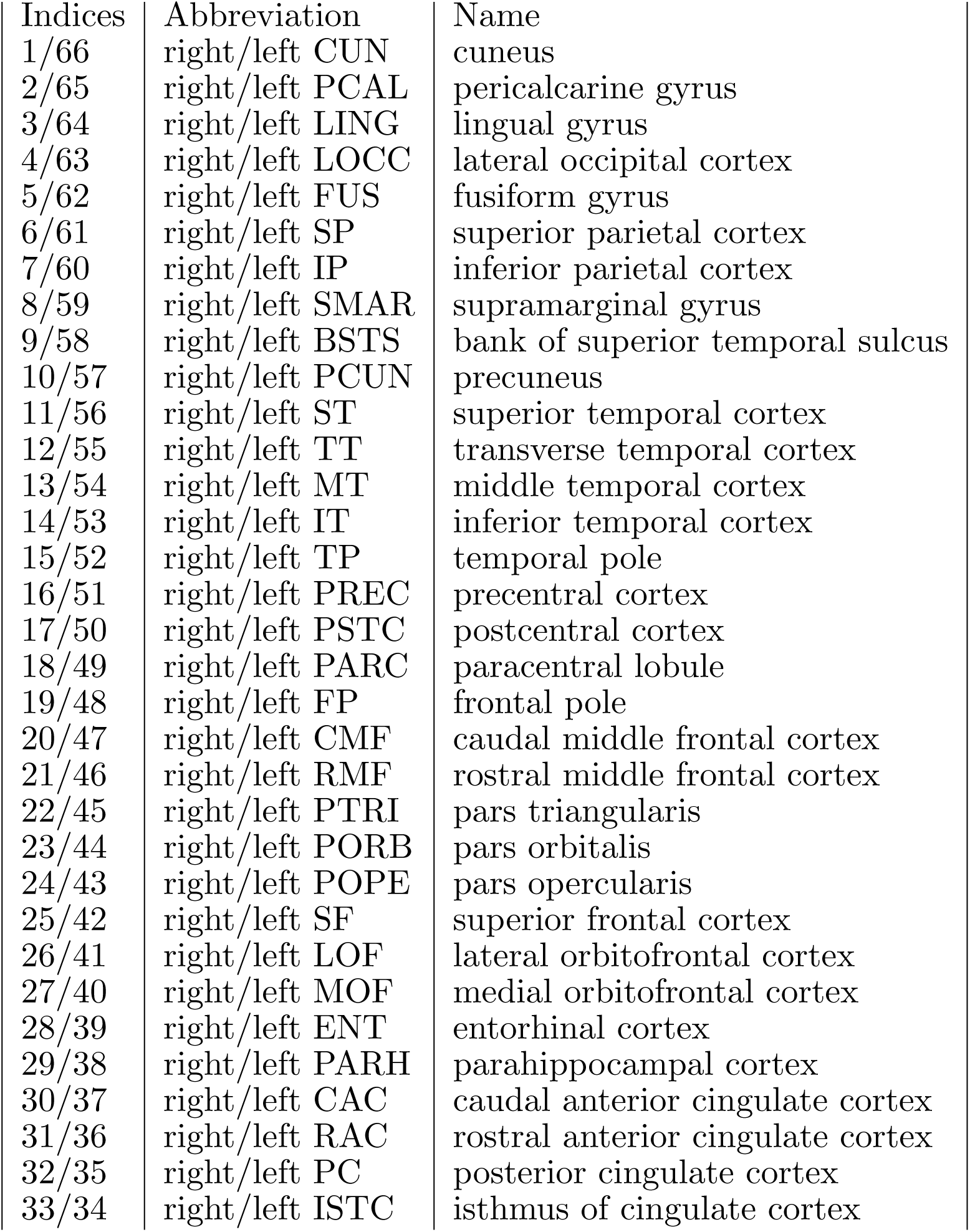
Table of ROI indices and labels.

| Indices | Abbreviation | Name |
| --- | --- | --- |
| 1/66 | right/left CUN | cuneus |
| 2/65 | right/left PCAL | pericalcarine gyrus |
| 3/64 | right/left LING | lingual gyrus |
| 4/63 | right/left LOCC | lateral occipital cortex |
| 5/62 | right/left FUS | fusiform gyrus |
| 6/61 | right/left SP | superior parietal cortex |
| 7/60 | right/left IP | inferior parietal cortex |
| 8/59 | right/left SMAR | supramarginal gyrus |
| 9/58 | right/left BSTS | bank of superior temporal sulcus |
| 10/57 | right/left PCUN | precuneus |
| 11/56 | right/left ST | superior temporal cortex |
| 12/55 | right/left TT | transverse temporal cortex |
| 13/54 | right/left MT | middle temporal cortex |
| 14/53 | right/left IT | inferior temporal cortex |
| 15/52 | right/left TP | temporal pole |
| 16/51 | right/left PREC | precentral cortex |
| 17/50 | right/left PSTC | postcentral cortex |
| 18/49 | right/left PARC | paracentral lobule |
| 19/48 | right/left FP | frontal pole |
| 20/47 | right/left CMF | caudal middle frontal cortex |
| 21/46 | right/left RMF | rostral middle frontal cortex |
| 22/45 | right/left PTRI | pars triangularis |
| 23/44 | right/left PORB | pars orbitalis |
| 24/43 | right/left POPE | pars opercularis |
| 25/42 | right/left SF | superior frontal cortex |
| 26/41 | right/left LOF | lateral orbitofrontal cortex |
| 27/40 | right/left MOF | medial orbitofrontal cortex |
| 28/39 | right/left ENT | entorhinal cortex |
| 29/38 | right/left PARH | parahippocampal cortex |
| 30/37 | right/left CAC | caudal anterior cingulate cortex |
| 31/36 | right/left RAC | rostral anterior cingulate cortex |
| 32/35 | right/left PC | posterior cingulate cortex |
| 33/34 | right/left ISTC | isthmus of cingulate cortex |

Another conceptual difference between the two approaches is the Gaussian assumption for the parameter distribution. For example, DCM relies in many cases on a single generative model with a distribution for each parameters —typically determined by a mean and a variance for each connection weights— and selects the model that provides the best evidence for all observed FC matrices. Instead, we estimate a single MOU-EC matrix per session, which builds a distribution of point samples for each connection. It remains to explore the implications of the difference in the natures of the estimate distributions and the detection of significant changes in MOU-EC —using the Bayesian machinery for DCM and standard machine learning in our case.

## 5 Network-oriented analysis for interpreting collective BOLD dynamics across conditions

So far, we have shown how MOU-EC estimates can be used for the classification of cognitive conditions using machine learning. Now we turn to the collective interpretation of MOU-EC links as a network. Tools from graph theory have been extensively applied to explore the brain network and understand how cognitive functions were distributed over subnetworks of ROIs. The study of both FC or SC has revealed a hierarchical organization of modular networks that are interconnected by hubs [85, 145, 157], which can be modulated depending on the subject’s condition [104, 13, 131].

A limitation in many SC and FC studies comes from the direct application of graph measures defined for binary data. In such cases SC or FC matrices are often binarized using an arbitrary threshold to obtain a graph, discarding the information conveyed by the graded values. For SC, this gives the skeleton of strong connections. Similar methods have also been used on FC [53, 38], but the use of a threshold to binarize FC seems more arbitrary, especially as many pairs of brain regions exhibit strong correlation. Another important aspect is often overlooked: brain connectivity measures such as FC are inferred from signals that have a temporal structure. This means that network analysis should take time into account. A possibility is to analyze the evolution of snapshot networks over time using dynamic FC based on sliding windows applied to the observed BOLD signals [94, 141, 6].

Going a step further, a particular focus has been on studying the relationship between the SC and FC [138, 15], in particular using dynamic models at the mesoscopic or macroscopic level [88, 63, 39, 113, 103, 20, 130, 56]. Following, graph analysis can be performed on the modeled BOLD activity [40, 55]. Nonetheless, it can be argued that such “integration” measures somehow rely on the observed phenomena (in empirical or model activity) rather than their causes. Rather, dynamic models enable the application of graph measures to the underlying connectivity that generates the dynamics, as was recently proposed [120, 67].

Here we follow this novel direction with our adaptation of graph-like measures for the network dynamics themselves [68, 67], whose main details are presented in the following; see also Appendix E for mathematical details. This section illustrates the following aspects of network-oriented analysis for MOU-EC:

- How to interpret collective changes in MOU-EC in their contribution to shape the network dynamics? We introduce dynamic communicability with the notion of ‘integration time’ that discriminates early and late interactions between ROIs in response to a (local) perturbation in the network via MOU-EC.
- Are network measures relevant for discriminating cognitive conditions? By doing so we want to obtain dynamics-specific biomarkers.
- Between the level of individual connections and the global network, are intermediate scales relevant to interpret the MOU-EC estimates? Can we detect functional communities of ROIs and are they task-specific?

### 5.1 Dynamic flow as graph-like measure for interactions between ROIs across ‘integration time’

Our definition of *dynamic flow* takes advantage of the information provided by the dynamic model, here applied to the MOU process [68]. As illustrated in Fig. 6A, MOU-EC quantifies the instantaneous causal interactions between pairs of ROIs and the Σ matrix measures the spontaneous or input activity level that each ROI receives (here represented only for ROI *i*). Built on those estimates, the dynamic flow measures the influence of a “perturbation” at a source ROI *i* onto a target ROI *k* after a given delay, which we call integration time. Importantly, it takes into account the network effect (or network response), meaning the contribution of indirect paths in the network (from ROI *i* to ROI *k* via ROI *j* in the figure). See Appendix E for the mathematical formulation based on the network impulse response (or Green function), which is analytically tractable for the MOU process.

**Figure 6:**
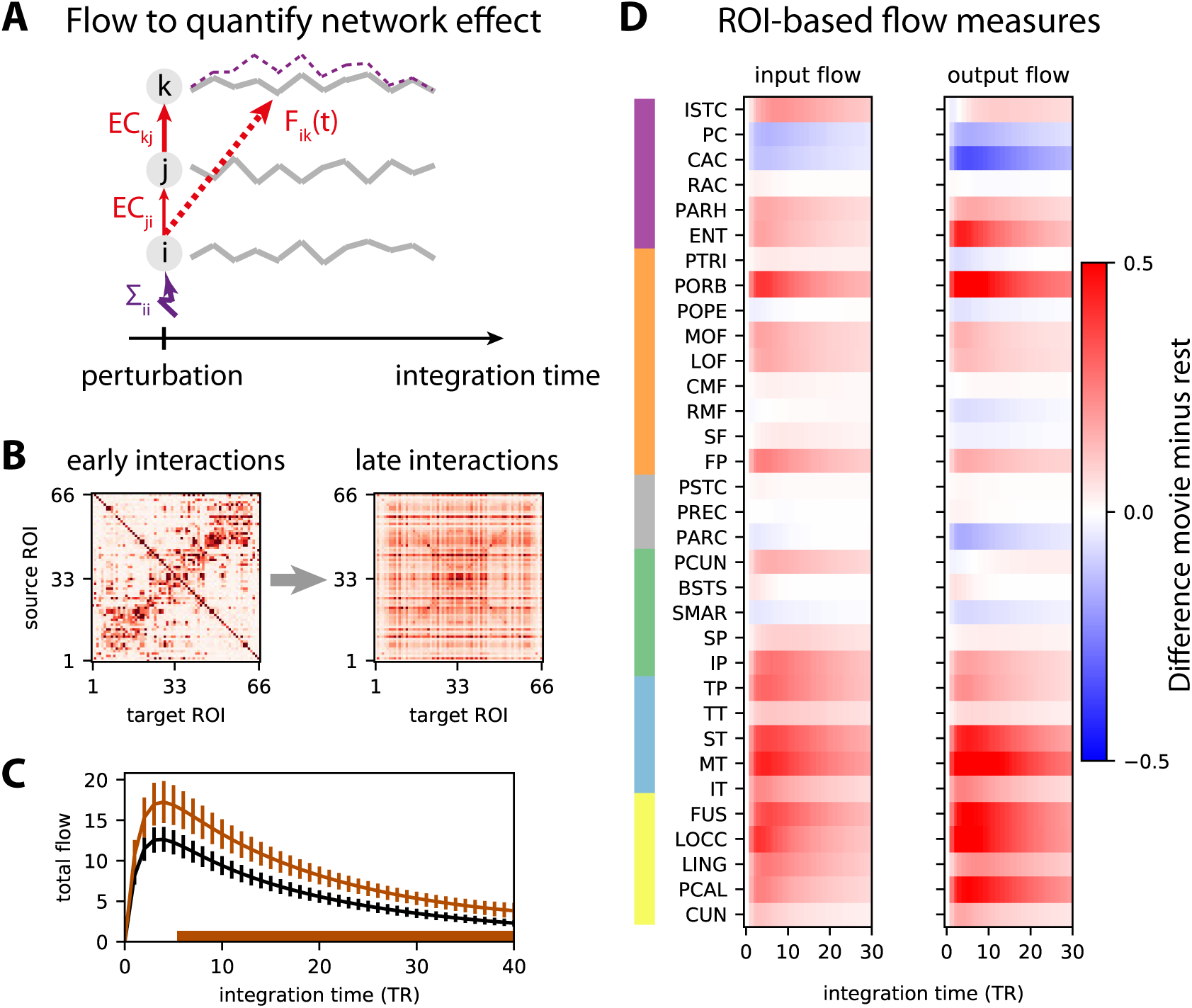
Network-oriented analysis of MOU-EC using dynamic flow. **A:** Conceptual illustration of dynamic communicability and flow. The propagation of extra activity resulting from a perturbation (here applied to ROI *i* in purple) develops over the ‘integration time’ and then fades away. The dashed purple curve represents the corresponding activity increase for ROI *k* (for legibility purpose, the equivalent for ROI *j* is not displayed). This defines interactions between ROIs that are not directly connected, like here from ROI *i* to ROI *k*. The dynamic communicability corresponds to interactions determined by the connectivity MOU-EC only, while the dynamic flow incorporates the amplitudes of ongoing perturbations that is quantified by Σ in the model. **B:** Two matrices representing the pairwise interactions as measured by dynamic communicability for all 66 ROIs at 2 integration times, in a single subject. Initially similar to MOU-EC values, the communicability becomes homogeneous. **C:** Total flow for rest and movie, which is the sum of the interactions between all pairs of ROIs (i.e. sum of all matrix elements in panel B) at each integration time. The error bars indicate the standard error of the mean over the subjects. The bottom brown horizontal line indicates where the total flow is larger for movie than for rest with *p <* 0.01 in a Mann-Whitney test for each integration time step (without correction). **D:** Change in input and output flow as a function of integration time, which sum all incoming and outgoing flow interactions for each ROI, respectively. The ROIs are ordered by anatomical areas and grouped in pairs of left-and right-hemisphere ROIs. The ROIs are anatomically grouped in the same manner as in Fig. 5A-B: yellow = occipital, cyan = temporal, green = parietal, gray = central, orange = frontal and purple = cingulate from bottom to top.

The dynamic flow thus aims to capture the dynamic response of the brain to local spontaneous activity (or external inputs) as it propagates from region to region in the network. A special case of the dynamic flow —termed *dynamic communicability* — that only depends on the MOU-EC (and not Σ) can also be defined assuming that all ROIs have independent spontaneous activity with the same intensity; it corresponds to a diagonal and homogeneous matrix Σ. Like the flow, dynamic communicability describes directional interactions, which differs from the symmetric interactions obtained from the direct application of graph theory to FC or SC [67]. It is worth stressing that the dynamic flow and communicability are graph-like measures that genuinely capture the properties of weighted networks, similar to the previously-proposed graph communicability [51, 50].

Considering again the fMRI dataset in Appendix A, we examine the dynamic flow at various scales in the brain network for the two conditions, movie and rest. This complements our previous analysis of the same dataset that showed that the modulations of MOU-EC between rest and movie implemented a selection of pathways in the cortex [66]. In addition, the stimulus load related to sensory inputs in the movie condition corresponded to an increase in Σ in the model for occipital and temporal ROIs. Recall that Σ quantifies the fluctuating BOLD activity intrinsic to each ROI that subsequently propagates via MOU-EC in the model. This motivates the use of the dynamic flow that incorporates Σ in order to characterize the whole-brain BOLD propagation, as compared to dynamic communicability. Note that the ‘effective drive’ defined in our previous study [66] corresponds to the flow at early integration time.

The sum of flow interactions provides insight about how much activity propagates throughout the whole cortical network, which is higher for movie compared to rest in Fig. 6C. Analyzing the interactions at the ROI level, the flow changes in Fig. 6D indicate differentiated effects among the ROIs, with increased/decreased listening for input flow and broadcasting for output flow. Here we observe most of the strongest increases in the output flow for occipital and temporal ROIs, which makes sense since the subjects both watch and listen the movie. A previous analysis based on MOU-EC [132] showed task-dependent changes for the outgoing connections from the rich club of ROIs that have dense anatomical connectivity, namely the precuneus (PCUN), superior parietal cortex (SP) and superior frontal cortex (SF), in agreement with previous results about the default-mode network also involving the post-cingulate gyrus (PC) [149, 150]. Here we only find a slight increase of input flow for PCUN and SP, which may be explained by the fact that the task is passive viewing and listening.

### 5.2 Integration in the brain network

Following experiments on the propagation of stimulus information from sensory areas to “high-level” brain areas in relation with being conscious of the stimuli [43, 28], the concept of integration was defined in models by quantifying the influence between regions while being in a broader network.

Following Fig. 6D, we use the dynamic flow to quantify how the BOLD activity propagates from the occipital and temporal ROIs, which exhibit an increased visual and auditory stimulus load in movie, to the rest of the brain. We thus consider the anatomical ROI groups in Fig. 7A (same as Fig. 5), where the stimulus load is represented by the purple perturbation applied to OCC and TMP [66]. Fig. 7B compares the summed flow between the ROI groups for rest and movie (in black and brown as in Fig. 6C). This reveals preferred pathways to the parietal and frontal ROIs (PAR and FRNT) when sensory information is integrated, as represented by the thick arrows in Fig. 7A.

**Figure 7:**
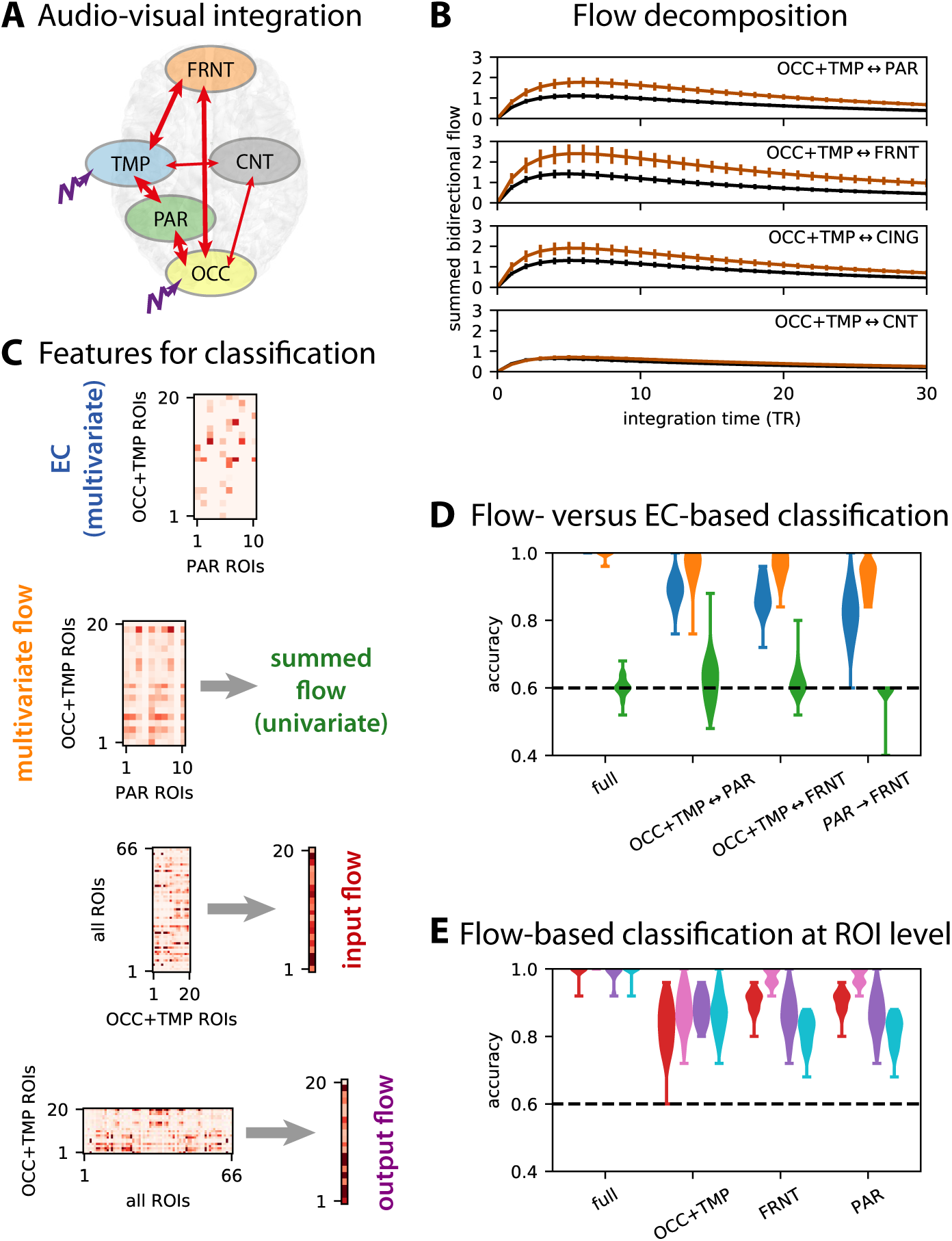
Integration measured by dynamic flow. **A:** Example subset of anatomical ROI groups: occipital (OCC), temporal (TMP), parietal (PAR), central (CNT) and frontal (FRNT). **B:** Comparison between the flow between the rest (in black) and movie (in brown) conditions. Similar to Fig. 6C, the curves are averages over subjects and corresponding sessions with error bars indicating the standard error of the mean. Each row corresponds to the flow summed over all ROIs of the indicated groups (bidirectionally), see also panel A where thick arrows correspond to strong increase in flow for the movie condition. **C:** Example of construction of features for the following classification. At the top, the EC matrix corresponds to connections for ROIs in groups and gives multivariate features when vectorized). Below, the flow matrix for the same ROIs equivalently gives multivariate features or can also be summed to obtain a univariate feature. Moreover, the flow matrix can be summed over columns or rows to obtain the input and output flows, here taking all 66 ROIs as sources (y-axis of the matrix) or targets (x-axis). **D:** Flow-based classification to discriminate rest versus movie in multivariate (orange violin plots) and univariate (green), compared to EC-based classification (blue). The x-axis indicates the ROI subgroups for which the flow interactions are selected. The dashed black lines indicate chance level, at 60% as in Fig. 4B. **E:** Same as panel D for the input flow (in red) and output flow (purple), which are compared to the input and output EC strengths (pink and cyan, respectively).

The machine-learning equivalent of the statistical testing for the summed flow between subnetworks in Fig. 7B gives the green violin plots in Fig. 7D. Here a good accuracy means that the bidirectional flow summed between all considered ROIs is strongly modulated between rest and movie. The fact that the results are close to chance level indicates strong variability in the overall flow increase across subjects. A different question, more related to biomarkers, is to which extent the detailed pattern of flow (multivariate flow, in orange) vary between movie and rest. This yields robust classification, which strikingly differs from the univariate (summed) flow. This indicates that the modification of interactions between ROIs is diverse beyond a simple global increase, hinting at the selection of specific pathways. The flow-based classification even outperforms MOU-EC (in blue) for the same matrix elements when focusing on particular ROI groups, see Fig. 7C for details. The discrepancy between the flow and MOU-EC shows the importance of network effects, for all three bidirectional pathways in Fig. 7D. The same classification can be performed with the input and output flows, see Fig. 7E. For interpretation purpose, the better performance for incoming than outgoing MOU-EC (pink versus cyan) suggests that ROIs change their pattern of listening/broadcasting communication, especially for PAR and FRNT. However, the overall effect is less pronounced when considering the dynamic flow that incorporates network effects (red versus purple). These results show that dynamics-based biomarkers that describe the propagation of BOLD activity can be built in the same way as connectivity-based biomarkers for the discrimination of cognitive tasks. Together with the distributed signature found in Fig. 5A, they also highlight the need for a whole-brain approach to understand the cortical reconfiguration [14], contrasting with previous studies relying on hypothesis testing for a-priori selected ROIs [72, 35, 30, 49].

### 5.3 Detection of functional communities determined by dynamic flow

In parallel to integration, the concept of complexity has been often used to differentiate brain states [143, 142, 7, 156]. Intuitively, complexity measures quantify the diversity of organizational motifs in a network, in the range of interest between the trivial extremes of full disconnectivity and full connectivity. A proxy for complexity is the detection of ROI communities because modular networks are a stereotypical model for complex networks, as was observed with SC [157]. Here we use the dynamic flow as a basis for the functional ROI communities in a data-driven manner, which contrasts with the anatomical grouping in Fig. 7A. Such methods developed initially for graphs with a metric called modularity based on weights [110]. Here we consider communities determined by the flow, namely ROIs with strong exchange of activity in the estimated dynamics [68].

Fig. 8A displays the evolution of communities over integration time for rest and movie. Strong coparticipation values indicate the robustness of the community structure across subjects. It shows a merging of almost all ROIs that eventually exchange flow in an even manner. This agrees with our previous results for resting state only and a different parcellation [67]. This speaks to a transition from segregated processing (within each diagonal block of left matrices) to a global integration. The number of communities can be taken as another proxy for the complexity in the brain network and our approach provides a dynamical perspective about it. The community merging goes hand in hand with the homogenization of the flow matrix in Fig. 8B, as quantified by the diversity of the matrix elements in the flow matrix that decreases and eventually stabilizes [68]. In our data the diversity is slightly higher for movie compared to rest.

**Figure 8:**
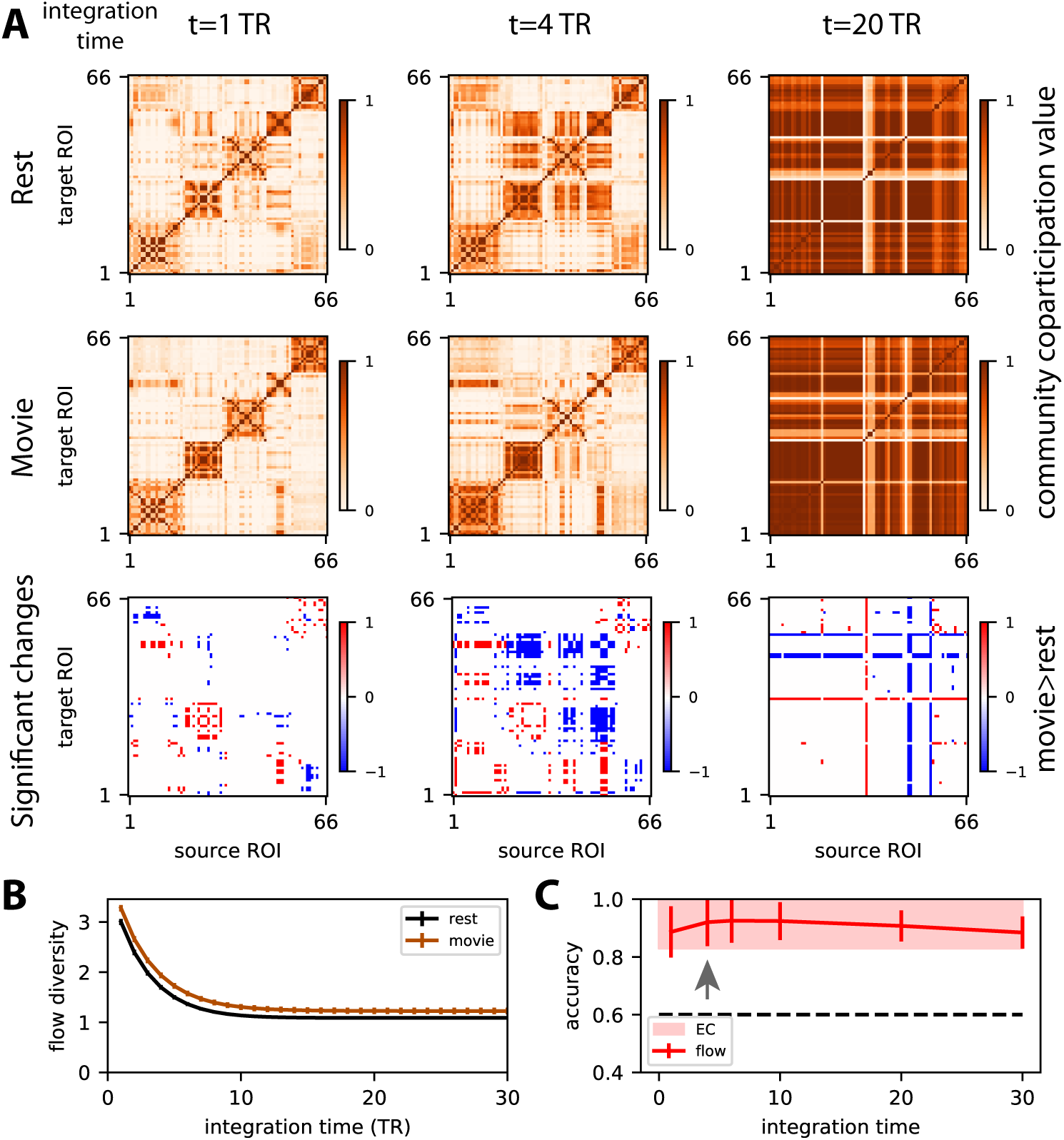
Community detection. **A:** Communities correspond to ROIs that bidirectionally exchange strong flow. Darker orange pixels indicate stable communities across the subjects for the each condition. The top row corresponds to rest and the middle row to movie for three integration times. The bottom row displays the significant increases (in red) and decreases (blue) of the community coparticipation values for movie with respect to rest. Here statistical testing is performed on each coparticipation value by using the Mann-Whitney test with *p* < 0.001 (without correction). The ROI ordering corresponds to the early community structure for rest. **B:** Flow diversity for rest (black curve) and movie (in brown) measuring the heterogeneity within each flow matrix when integration time increases; see Eq. (19). It is a proxy for the “complexity” of interactions between ROIs; for details see [68, 67]. **C:** Rest-versus-movie classification based on flow (whole-brain version of multivariate flow in Fig. 7C) as a function of integration time and comparison with EC-based classification. Starting *t* = 4 TRs indicated by the arrow, the performance reaches a maximum plateau. Here the train test consists of 20% of the samples and the test set 80% (the converse compared to before) in order to capture the strongest changes between rest and movie with classification. This exaggerates the evolution over integration time of the performance, which is lower compared to the previous plots.

Going a step further, we can examine the difference between the community structures of the two conditions, see the bottom row in Fig. 8A. This shows that the changes in dynamic flow for specific pairs of ROIs between rest and movie (Fig. 7B, D and E) have an important collective impact. The strongest differences appear when the network effects are very strong at *t* = 4 TRs. This is confirmed by Fig. 8C, where classification shows that the dynamic flow is most different between rest and movie at integration times corresponding to strong network effects (the arrow indicates the start of the plateau for maximum performance). These results speak to an intermediate organization of groups of ROIs between the local and global scales, which can be captured using our network-oriented analysis of MOU-EC.

## 6 Conclusions

This article illustrates how a model-based approach to whole-brain fMRI analysis combines the desired properties of predictability and interpretability. This goes beyond the goodness of fit for fMRI signals that is commonly used to evaluate and compare generative models. To do so, our framework links tools and concepts from dynamic systems, machine learning and network theory. This results in a consistent pipeline with controlled hypotheses because we use the same whole-brain dynamic model from the estimation to the analysis.

The analysis tools have been organized in a package written in the open-source language Python: https://github.com/mb-BCA/pyMOU for the MOU-EC estimation and https://github.com/mb-BCA/NetDynFlow for dynamic communicability and flow. The code to reproduce important figures using the data is available on https://github.com/mb-BCA/notebooks_review2019. The classification uses the library scikit-learn [2].

### General key points for analysis of multivariate neuroimaging data and corresponding results using MOU-EC

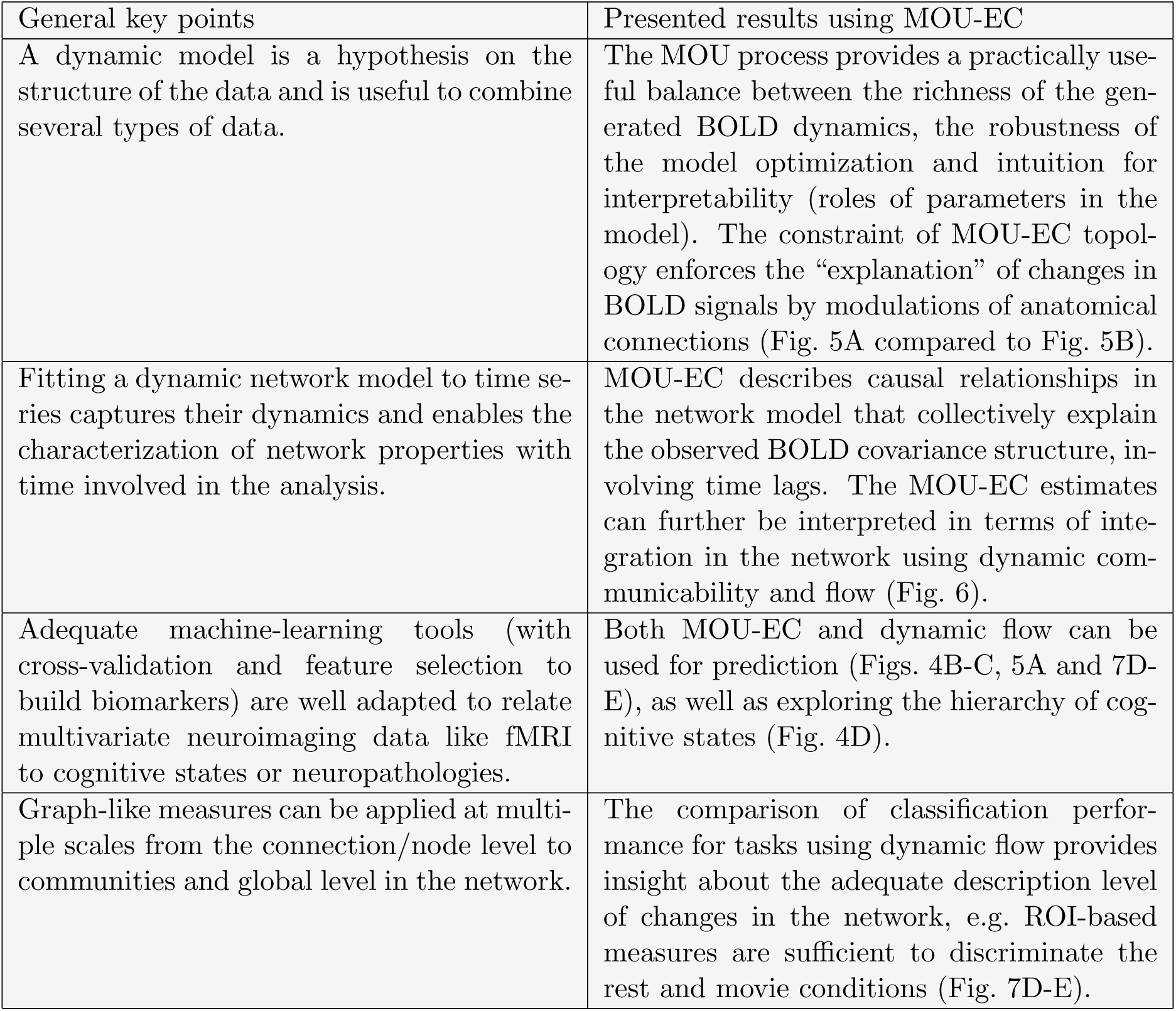

**General key points for analysis of multivariate neuroimaging data and corresponding results using MOU-EC:**
| General key points | Presented results using MOU-EC |
| --- | --- |
| A dynamic model is a hypothesis on the structure of the data and is useful to combine several types of data. | The MOU process provides a practically useful balance between the richness of the generated BOLD dynamics, the robustness of the model optimization and intuition for interpretability (roles of parameters in the model). The constraint of MOU-EC topology enforces the “explanation” of changes in BOLD signals by modulations of anatomical connections (Fig. 5A compared to Fig. 5B). |
| Fitting a dynamic network model to time series captures their dynamics and enables the characterization of network properties with time involved in the analysis. | MOU-EC describes causal relationships in the network model that collectively explain the observed BOLD covariance structure, involving time lags. The MOU-EC estimates can further be interpreted in terms of integration in the network using dynamic communicability and flow (Fig. 6). |
| Adequate machine-learning tools (with cross-validation and feature selection to build biomarkers) are well adapted to relate multivariate neuroimaging data like fMRI to cognitive states or neuropathologies. | Both MOU-EC and dynamic flow can be used for prediction (Figs. 4B-C, 5A and 7D-E), as well as exploring the hierarchy of cognitive states (Fig. 4D). |
| Graph-like measures can be applied at multiple scales from the connection/node level to communities and global level in the network. | The comparison of classification performance for tasks using dynamic flow provides insight about the adequate description level of changes in the network, e.g. ROI-based measures are sufficient to discriminate the rest and movie conditions (Fig. 7D-E). |

### 6.1 Spatio-temporal BOLD structure

By fitting the dynamic model to the spatio-temporal covariance structure of the BOLD signals, our approach captures the BOLD dynamics averaged over each fMRI session. The estimated MOU-EC can be seen as a “projection” of the BOLD signals on anatomically-constrained connectivity, thereby characterizing the dynamical state of the brain. In contrast, analyses based on the static FC can be related to a linear Gaussian model, and thus does not take the temporal dimension into account (i.e. BOLD values are considered as i.i.d. variables). Dynamic FC methods have been developed to incorporate time in the analysis [113, 117, 74, 25], but they do not genuinely capture the propagating nature of BOLD signals. Dynamic causal models have just been adapted to deal with whole-brain fMRI data [120, 114, 56] and it will be interesting to compare the estimated connectivities. It is also still unclear how short sessions or observation windows can be to robustly characterize dynamical states when calculating EC estimates or MAR-like coupling matrices in HMMs [151, 18], in comparison to dynamic FC methods that are typically used with windows of 30 to 60-TRs.

### 6.2 Biomarkers for cognition and neuropathologies

The robust estimation of the (multivariate) parameters allows for a powerful representation of cognitive states by the MOU-EC estimates (Fig. 4), as well as the extraction of task-specific biomarkers (Fig. 5). An important point considering connectivity measures is that machine learning solves the problem of multiple comparisons: instead of determining p-values and then a threshold for connections with significant changes, we can characterize which MOU-EC links —as features— contribute to the correct classification. It has been recently advocated that machine learning can be used to avoid “excessive reductionism”, namely linking brain activity in localized regions to narrow experimental protocols in a way that cannot generalize to more complex protocols [146]. Although the focus has been on machine-learning techniques, we remind that hypothesis testing of significant changes for preselected ROIs or links can be performed in our framework (see Fig. 7B with the dynamic flow). In that case, whole-brain changes can also provide a baseline reference for the considered links or ROIs.

On the technical side, it remains to be explored how the classification performance is affected by the resolution of various parcellations [33, 47, 71], as was recently done with FC-like connectivity measures [34]. The trade-off to solve is between the richness of the connectivity measure (more links can reproduce richer environments) and their robustness (more parameters may result in noisier estimation). The alignment of distinct datasets with various MRI protocols also remains a difficult question for practical application to large datasets [148].

In parallel to the study of cognition, the presented framework can be applied to neuropathologies with the aim to inform clinical diagnosis [101]. The rationale is that BOLD activity specifically reflects neuropathologies, even in the resting state [77, 87]. If SC is increasingly used for strokes or Alzheimer disease that strongly impact the brain anatomy, fMRI may provide additional information [134, 79]. Other diseases like depression [46] or some motor disorders [123] may be better investigated using functional imaging. Importantly, many neural diseases are likely to affect multiple brain regions and thus require adequate tools to capture their distributed nature. Another direction where machine learning is a key tool is the development of personalized medicine, adapting the models to individual patients beyond group analyses [155].

### 6.3 Network-oriented analysis of dynamics

The network analysis for our dynamic model relies on the dynamic flow, a graph-like connectivity measure that describes the integration of propagating activity at multiple scales in the network (Figs. 7 and 8). In particular, small changes in EC that are not individually significant can collectively induce large changes in the dynamic flow, especially when feedback loops are involved. In this sense, the dynamic flow quantitatively captures the interplay between parameters, so the resulting communication cannot always be simply understood at the level of single estimated parameters. As an example, many small changes in EC (below significance level) may collectively result in a strong dynamical effect, as seen for task A in Fig. 9. Strong coordinated changes in EC result in large changes in the dynamics (task B). Moreover, changes in input properties may only be captured by the dynamics, in particular when the connectivity does not change (task C). Building biomarkers that capture network effects is important to make use of the multivariate nature of the fMRI data. This is important when interpreting data in terms of concepts such as integration, segregation and complexity [142, 43, 42, 156]. An interesting direction for future work is the study of directional properties of the flow, especially in the characterization fo functional communities.

**Figure 9:**
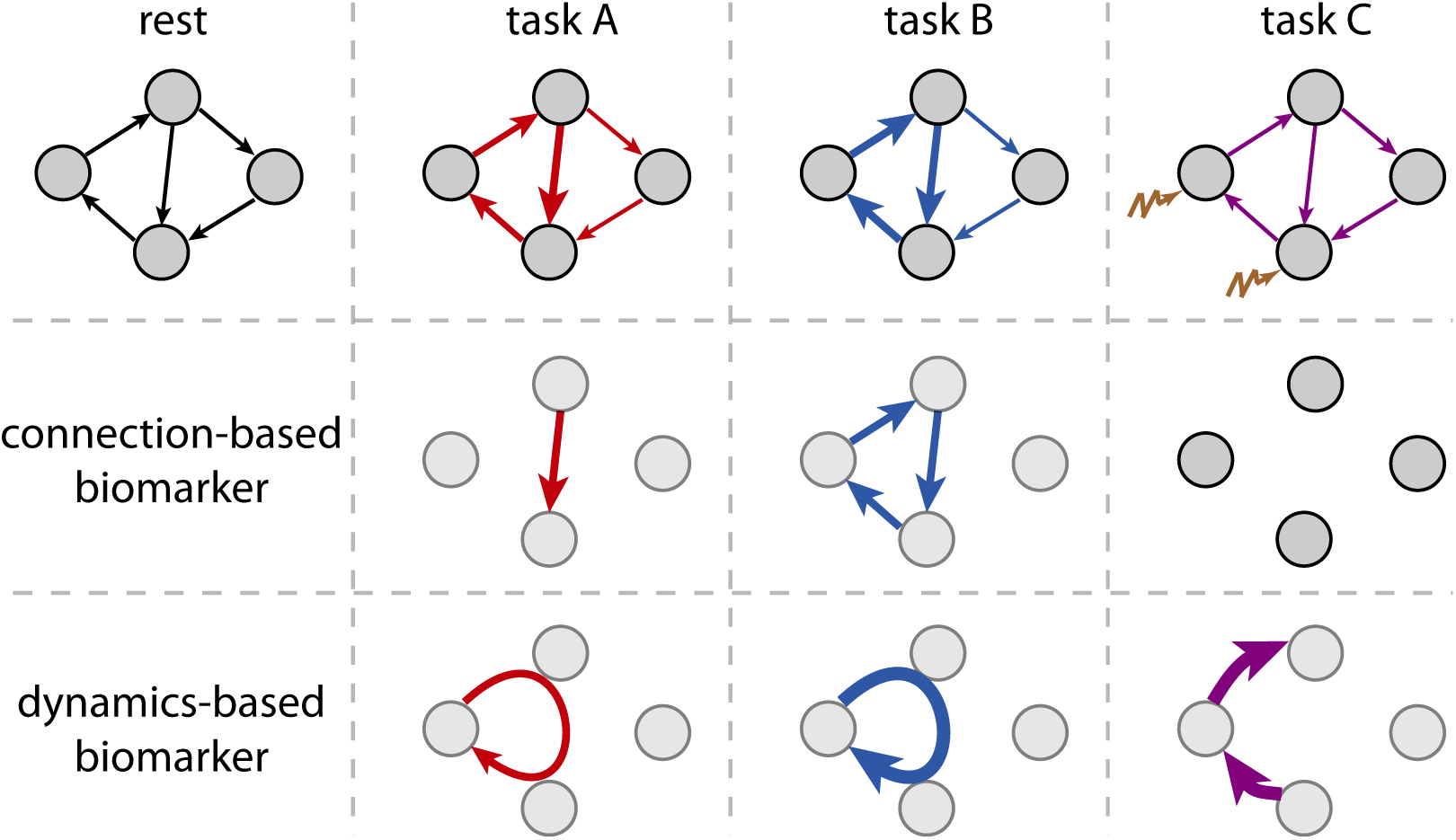
Schematic representation of the extraction of biomarkers for three tasks compared to rest. Connection-wise biomarkers corresponds to changes in EC while the bottom row refers to biomarkers for changes in dynamic flow. Tasks A and B correspond to two modulations of the connectivity, which lead to similar changes in the dynamic flow (stronger for B). For task C, extra inputs are received by two nodes, which increase the dynamic flow towards the corresponding two target nodes.

### 6.4 Limitations and future work

As a first warning, we remind that any interpretation of BOLD in term of brain communication relies on the assumption that changes in neuronal activity are reliably reflected in the BOLD signals, which is still under debate [73, 100, 84]. In practice, the preprocessing of fMRI signals must strive to increase the signal-to-noise ratio [116, 115].

Recall also that causal relationships between brain regions come from the directed connectivity in the model, whose “best” parametrization is estimated from the data. The estimation procedure is crucial, in particular for the dynamic flow here since the asymmetry of MOU-EC strongly affects the resulting activity propagation in the model. Moreover, without a proper experimental interaction with the brain activity like using stimulation, the notion of causality is subject to the initial model assumption [93, 89]. This remark also applies to other methods for directed connectivity, such as Granger causality and DCM [144].

A key distinction of our approach compared to DCM is that our MOU-EC (or FC) estimates are calculated for each fMRI session, so we obtain a sample distribution for connection weight over all subjects for each condition (e.g. task). In contrast, DCM usually considers a single model per condition that aims to represent the variability over subjects with a parametric distribution (mean and variance) for each connection weight. The limitations and advantages of both approaches in terms of statistical power and generalizability remain to be practically compared, especially for large datasets.

Regarding the hemodynamics and explicit HRF modeling discussed at the end of the section presenting the MOU model, a quantitative comparison between MOU-EC, DCM and Granger causality analysis to extract biomarkers or interpret fMRI data in terms of neuronal coupling is necessary in the future. A particular point concerns the observed heterogeneity of HRF across different brain regions and how it affects the estimated connectivity and its directionality [81]. Previous works also suggest that the HRF is important for the reliability of connectivity analysis [70, 111]. This may increase the number of parameters in the model, so the robustness of the estimation should be verified using both forward modeling in synthetic whole-brain networks [135] and with classification from real data with known categories (e.g. subjects with several sessions each). A first step could be to extend the MOU-EC as a state-space model with a first-order filter for HRF [127].

## Appendix

The following sections summarize details about the data and mathematical framework used in this article. For machine learning, the readers are referred to the online documentation http://scikit-learn.org/ of the scikit-learn library [2].

### A Movie dataset

The experimental procedure was approved by the Ethics Committee of the Chieti University and the participants signed a written informed consent. We use the data from 22 right-handed young and healthy volunteers (15 females, 20-31 years old) that had recordings for both a resting state with eyes opened and a natural viewing condition (30 minutes of the movie ‘The Good, the Bad and the Ugly’).

The fMRI time resolution (TR) is 2 seconds and each session of 10 minutes corresponds to 300 frames (or volumes). The preprocessing performed using SPM8 involves the coregistration of structural and functional MRI, spatial normalization in the MNI coordinates and motion correction. The data are parcellated in *N* = 66 ROIs [80]. The ROIs are listed in Table 1, indexed in the order of Figs. 5 and 6. The generic structural matrix is obtained from averaging individual matrices obtained using the diffusion spectrum imaging. A threshold was then applied to retain the 27% largest connections. Inter-hemispheric connections between homotopic ROIs were added, increasing the density to 28%. The same generic SC matrix is used to determine the MOU-EC topology for all subjects.

We refer the readers to previous publications [86, 99, 66] for further details.

### B Functional connectivity measures

For each session of *T* = 300 time points (2 for rest and 3 for movie), we denote the BOLD time series by 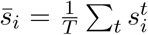 for each region 1 ≤ *i* ≤ *N* with time indexed by 1 ≤ *t* ≤ *T*. The mean signal is 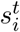 for all *i*. Following [69], the empirical FC consists of two BOLD covariance matrices (see the blue and green matrices in the top box of Fig. S1), without and with time lag:

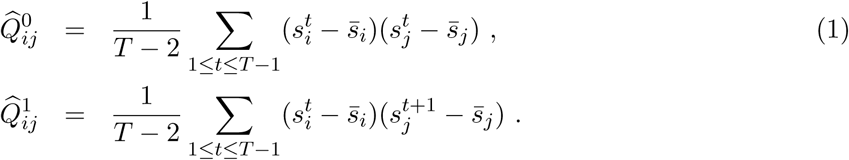

Pearson correlation can be calculated from the covariances in Eq. (1):

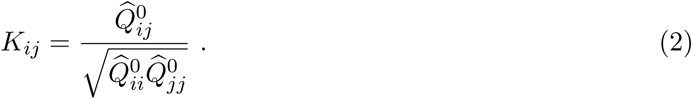

### C Multivariate Ornstein-Uhlenbeck (MOU) process to model whole-brain dynamics

The network dynamics are determined by two sets of parameters:

- the time constant *τ*_*x*_ is an abstraction of the
- the network effective connectivity (MOU-EC) between these ROIs embodied by the matrix *C*, whose topological skeleton is determined by structural data;
- the local variability embodied in the matrix Σ inputed individually to each of the *N* = 66 ROIs.

**Figure S1:**
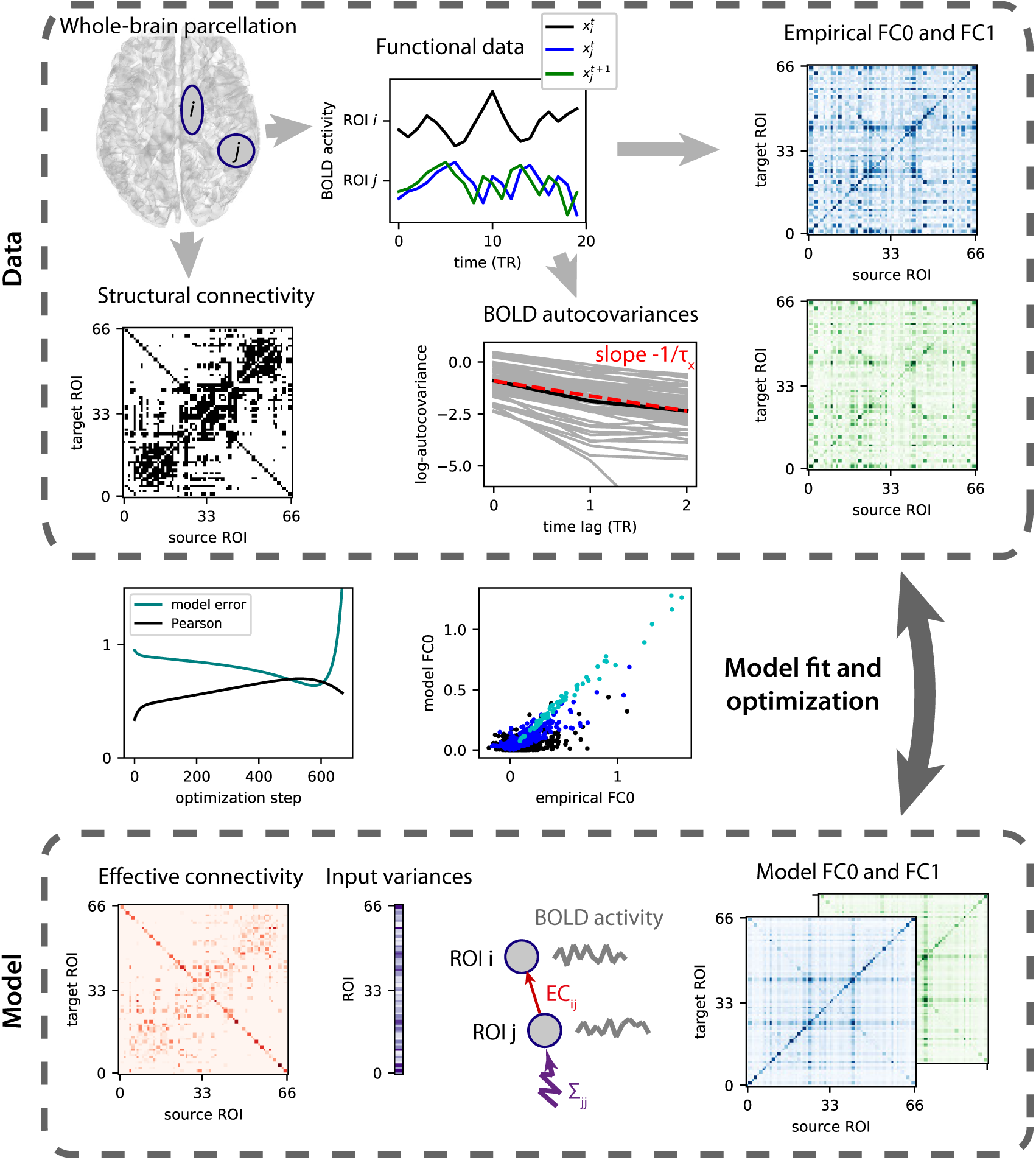
Dynamic model and optimization procedure to capture BOLD dynamics. From the BOLD signals of the parcellation (top left in top box), two empirical FC matrices (blue and green matrices) are calculated, both without and with time lag as depicted by the blue and green curve. The dynamic network model describes the activity of the ROIs (only two are represented by the grey circles) and comprises a parameter for the spontaneous activity of each ROI (Σ in the bottom box) and for each directed connection between ROIs (effective connectivity, MOU-EC). The topology of existing connections is determined by the structural connectivity (black matrix in top box). The optimization of the model is similar to a gradient descent during which the model FC matrices are evaluated at each step and compared to their empirical counterparts. From the differences between the corresponding matrices an update of the MOU-EC and Σ parameters is calculated and the procedure is repeated until reaching a minimum for the model error (light gray-blue curves in the left central panel between the boxes), which corresponds to a high value for the Pearson correlation between the matrix elements of the model and empirical FCs (black curve). Before the optimization, the time constant *τ*_*x*_ of the BOLD autocovariances (see log-plot) is evaluated to calibrate the nodal dynamics in the model. Details can be found in [69, 66].

The activity variables obey are described by a multivariate Ornstein-Uhlenbeck process. Each activity variable *x*_*i*_ of node *i* decays exponentially according to the time constant *τ*_*x*_ and evolves depending on the activity of other populations:

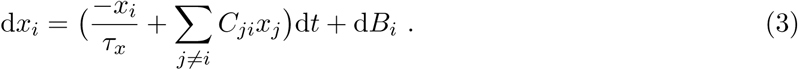

Compared to our previous work [69, 66], we use a different convention for the weights *C*_*ji*_ from ROI *j* to ROI *i*, in line with graph theory. In matrix notation it now reads

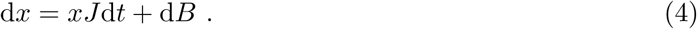

Here the fluctuating inputs d*B*_*i*_ are independent and correspond to a diagonal covariance matrix Σ, as represented in purple by the vector of variances Σ_*ii*_ in the bottom box of Fig. S1. In the model, all variables *x*_*i*_ have zero mean. Their spatiotemporal covariances are denoted by 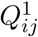, where *τ* ∈ {0, 1} is the time lag, and correspond to the the blue and green matrices in the bottom box of Fig. S1. They can be calculated by solving the consistency equations:

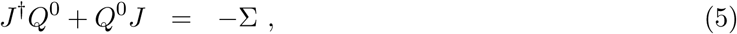

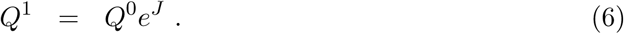

The first equation is the continuous Lyapunov equation, which can be solved using the Bartels-Stewart algorithm in scipy [1]. Here *J* is the Jacobian of the dynamical system and depends on the time constant *τ*_*x*_ and the network MOU-EC: 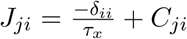, where *δ*_*ji*_ is the Kronecker delta and the superscript † denotes the matrix transpose. In the second equation *e* denotes the matrix exponential.

Ideally, the model should be extended to incorporate subcortical areas and the relevance of input cross-correlations inputs should be evaluated for all ROI pairs [66].

### D Parameter estimation procedures

#### D.1 Lyapunov optimization or natural gradient descent

We tune the model such that its covariance matrices *Q*^0^ and *Q*^1^ reproduce the empirical FC0 and FC1 matrices in Fig. 2B, denoted here by 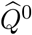 and 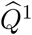, respectively. The uniqueness of this maximum-likelihood estimation follows from the bijective mapping from the model parameters *τ*_*x*_, *C* and Σ to the FC pair (*Q*^0^, *Q*^1^). The iterative optimization procedure for *C* is similar to the original version [69], which can be related to the concept of “natural” gradient descent [4] that takes the non-linearity of the mapping between *J* and the matrix pair *Q*^0^ and *Q*^1^ in the second line of Eq. (5). Note that the parameters *τ*_*x*_ and Σ also follow a gradient descent now.

For each session (subject and condition), we calculate the time constant *τ*_ac_ associated with the exponential decay of the autocovariance averaged over all ROIs using time lags 0 and 1 TR:

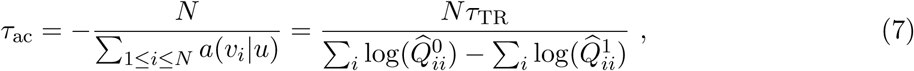

where *a*(*v*_*i*_|*u*) is the slope of the linear regression of 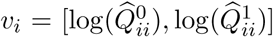 by *u* = [0, 1], and *τ*_TR_is the value of the TR in seconds. Note that one can also use more time lags —e.g. up to 2 TRs— to assess the stability of the estimated *τ*_ac_, as was previously analyzed for resting-state data [69]. The model is initialized with *τ*_*x*_ = *τ*_ac_ and no connectivity *C* = 0, as well as unit variances without covariances (Σ_*ij*_ = *δ*_*ij*_). At each step, the Jacobian *J* is straightforwardly calculated from the current values of *τ*_*x*_ and *C*. Using the current Σ, the model FC matrices *Q*^0^ and *Q*^1^ are then computed from the consistency equations, using the Bartels-Stewart algorithm to solve the Lyapunov equation.

The difference matrices 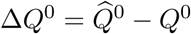 and 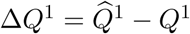 determine the model error

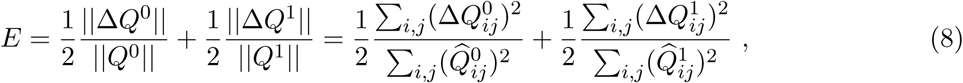

where each term - for FC0 and FC1 - is the matrix distance between the model and data covariances, normalized by the latter.

The following Jacobian update can be applied to decrease the model error *E* at each optimization step similar to a gradient descent:

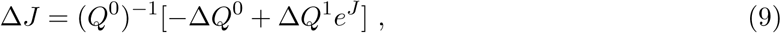

It turns out that a modified update is more robust to empirical noise in practice

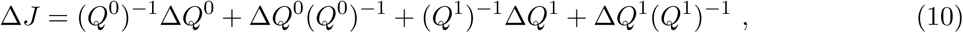

which would corresponds to a proxy error based on the matrix logarithm:

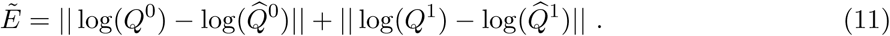

See [91] for a comparison of optimization methods. From the Jacobian update Δ*J*, we obtain the connectivity update:

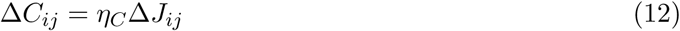

for existing connections only; other weights are forced at 0. We also impose non-negativity for the MOU-EC values during the optimization.

To take properly the effect of cross-correlated inputs into account, the Σ update has been adjusted from the heuristic update in the first study [69] to a gradient descent [66]:

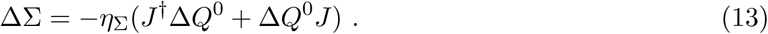

As with weights for non-existing connections, only diagonal elements of Σ —and possible cross-correlated inputs [66]— are tuned; others are kept equal to 0 at all times.

Last, to compensate for the increase of recurrent feedback due to the updated *C*, one can also tune the model time constant *τ*_*x*_ as

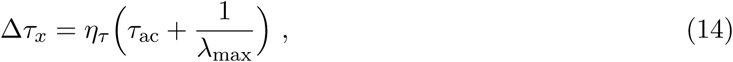

where *λ*_max_ is the maximum negative real part of the eigenvalues of the Jacobian *J*. The rationale is to avoid an explosion of the network dynamics (when *λ*_max_ → 0) while letting the model connectivity *C* develop a non-trivial structure to reproduce FC.

Repeating the parameter updates, the best fit corresponds to the minimum of the model error *E*.

#### D.2 Heuristic optimization

Instead of the derivation of the Jacobian in Eq. (10) that takes into account the nonlinearity of the mappings in Eq. (5), one can use a greedy algorithm to optimize the weights to fit an objective measure on the data. Here we update the weights for the model to reproduce the correlation matrix *K* in Eq. (2):

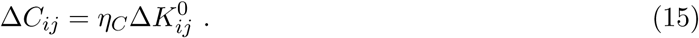

Note that only correlations 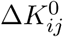 for existing connections *i* → *j* and *j* → *i* are taken into account in this update.

### E Dynamic communicability and flow for network analysis

Following our previous study [68], we firstly define *dynamic communicability* to characterize the network interactions due to the MOU-EC connectivity *C*, ignoring the input properties Σ. Our definition is adapted to study complex networks associated with realistic (stable) dynamics where time has a natural and concrete meaning. In comparison, a previous version of communicability for graphs [51] relied on abstract dynamics. The basis of our framework is the network response over time, or Green function, which is the basis of the concept of dynamic communicability, which focuses on the temporal evolution of such interactions. Although we focus on the MOU process here, our framework can be easily adapted to distinct local dynamics for which the Green function is known. In addition to the MOU-EC matrix *C*, the MOU dynamics is determined by the input properties, so we use the *dynamic flow* in the present article (Figs. 6 and 8).

#### E.1 Definitions

Formally, dynamic communicability is the “deformation” of the Green function *e*^*Jt*^ of the MOU process due to the presence of the (weighted and directed) matrix *C*, as compared to the Green function 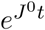 corresponding to the Jacobian with leakage only and no connectivity, 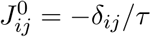. It is defined as the family of time-dependent matrices depicted in Fig. 6A:

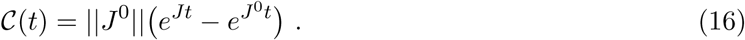

The scaling factor 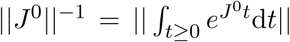 where ‖ · ‖ is the L1-norm for matrices (i.e., sum of elements in absolute value) is used for normalization purpose [68]. Recall that *t* ≥ 0 here is the time for the propagation of activity in the network, referred to as ‘impulse-response time’ in the figures.

To incorporate the effect of local spontaneous activity or excitability (inputs in the model), we define the dynamic flow that fully characterizes the complex network dynamics [68]. The input statistics of interest for a stable MOU process correspond to the input (co)variance matrix Σ, which are independent parameters from the MOU-EC matrix *C*. This is represented by the purple arrows in the left diagram of Fig. 6A, indicating that the fluctuation amplitude is individual for each ROI. The Σ matrix may be non-diagonal when ROIs experience cross-correlated noise [66], as represented by the purple dashed arrows. The dynamic flow describes the propagation of local fluctuating activity over time via the recurrent connectivity and is defined by the

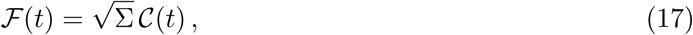

where 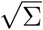 is the real symmetric “square-root” matrix of the input covariance matrix, satisfying 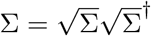. Dynamic communicability is thus a particular case of the flow for homogeneous input statistics.

From the time-dependent matrices ℱ(*t*), we define the total flow that sums all interactions

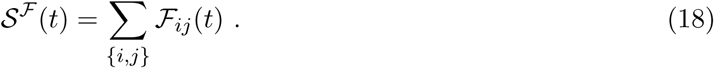

Total communicability for graphs has been used to define a version of centrality [12]. Here the proximity between ROIs correspond to how much flow they exchange. We also define the diversity (akin to heterogeneity) among the ROI interactions in the time-dependent matrices ℱ(*t*), which can be seen as a proxy for their homogenization over time:

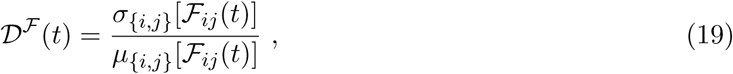

defined as a coefficient of variation where *µ*_*{i,j}*_ and *σ*_*{i,j}*_ are the mean and standard deviation over the matrix elements indexed by (*i, j*).

We also define the input and output flows for each node:

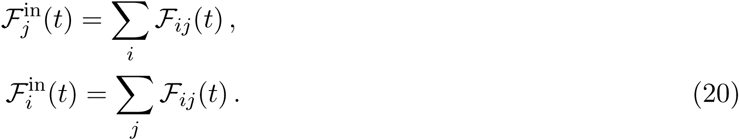

#### E.2 Community detection

To detect communities from ℱ(*t*) in Fig. 8D, we rely on Newman’s greedy algorithm for modularity [110] that was originally designed for weight-based communities in a graph. Adapting it here to the flow matrix ℱ(*t*) at a given time *t*, we seek for communities where ROIs have strong bidirectional flow interactions. In the same manner as with weighted modularity, we calculate a null model for MOU-EC:

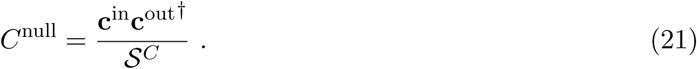

Note that we preserve the empty diagonal. The resulting matrix contains from the expected weight for each connection, given the observed input strengths **c**^in^ and output strengths **c**^out^; 𝒮^*C*^ is the total sum of the weights in *C*. Then we caclulate ℱ^null^(*t*) using Eq. (17) with *C*^null^ instead of *C*. Starting from a partition where each ROI is a singleton community, the algorithm iteratively aggregates ROIs to form a partition of *K* communities denoted by *S*_*k*_ that maximizes the following quality function:

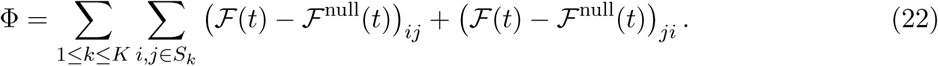

At each step of the greedy algorithm, the merging of two of the current communities that maximizes the increase of Φ is performed. Note that communicability-based communities can be defined similarly using C(*t*) and the corresponding null model C^null^(*t*).

## Notes

#### Summary of Updates

revised version and reference to new python package

